# Quantitative modeling links *in vivo* microstructural and macrofunctional organization of human and macaque insular cortex, and predicts cognitive control abilities

**DOI:** 10.1101/662601

**Authors:** Vinod Menon, Gallardo Guillermo, Mark A. Pinsk, Van-Dang Nguyen, Jing-Rebecca Li, Weidong Cai, Demian Wassermann

**Affiliations:** Department of Psychiatry & Behavioral Sciences, Stanford University School of Medicine Stanford, CA 94305; Department of Neurology & Neurological Sciences, Stanford University School of Medicine Stanford, CA 94305; Stanford Neurosciences Institute, Stanford University School of Medicine Stanford, CA 94305; Athena, Inria Sophia Antipolis, Université Côte d’Azur, 2004 route des Lucioles 06902 Sophia Antipolis CEDEX, France; Princeton Neuroscience Institute, Princeton University Princeton, NJ 08544; Department of Computational Science and Technology Royal Institute of Technology in Stockholm Stockholm, NJ 08544; Defi, Inria Saclay Île-de-France, École Polytechnique Université Paris Sud 1 Rue Honoré d’Estienne d’Orves 91120 Palaiseau, France; Parietal, Inria Saclay Île-de-France, CEA Université Paris Sud 1 Rue Honoré d’Estienne d’Orves 91120 Palaiseau, France

**Keywords:** insula, VEN, salience network, cognitive control, behavior, neural, circuits

## Abstract

The human insular cortex is a heterogenous brain structure which plays an integrative role in guiding behavior. The cytoarchitectonic organization of the human insula has been investigated over the last century using postmortem brains but there has been little progress in noninvasive *in vivo* mapping of its microstructure and large-scale functional circuitry. Quantitative modeling of multi-shell diffusion MRI (dMRI) data from 440 HCP participants revealed that human insula microstructure differs significantly across its functionally defined dorsal anterior, ventral anterior, and posterior insula subdivisions that serve distinct cognitive and affective functions. The microstructural organization of the insula was mirrored in its functionally interconnected circuits with the anterior cingulate cortex that anchor the salience network, a system important for adaptive switching of cognitive control systems. Novel validation of the human insula findings came from quantitative dMRI modeling in macaques which revealed microstructural features consistent with known primate insula cytoarchitecture. Theoretical analysis and computer simulations, using realistic 3-dimensional models of neuronal morphology from postmortem tissue, demonstrated that dMRI signals reflect the cellular organization of cortical gray matter, and that these signals are sensitive to cell size and the presence of large neurons such as the von Economo neurons. Crucially, insular microstructural features were linked to behavior and predicted individual differences in cognitive control ability. Our findings open new possibilities for probing psychiatric and neurological disorders impacted by insular cortex dysfunction, including autism, schizophrenia, and fronto-temporal dementia.

**Statement of Significance:** The human insular cortex is a heterogenous brain structure which plays an integrative role in identifying salient sensory, affective, and cognitive cues for guiding attention and behavior. It is also is one of the most widely activated brain regions in all of human neuroimaging research. Here we use novel quantitative tools with *in vivo* diffusion MRI in large group (N=440) of individuals to uncover several unique microstructural features of the human insula and its macrofunctional circuits. Crucially, microstructural properties of the insular cortex predicted human cognitive control abilities, in agreement with its crucial role in adaptive human behaviors. Our findings open new possibilities for probing psychiatric and neurological disorders impacted by insular dysfunction, including autism, schizophrenia, and fronto-temporal dementia.

---

The human insular cortex plays a critical role in identifying salient sensory, affective, and cognitive cues for guiding attention and behavior(1–4). Critically, it is also is one of the most widely activated brain regions in all of human neuroimaging research(5–7). Dysfunction of the human insula and its interconnected regions are now thought to be core features of many psychiatric and neurological disorders(8, 9). However, little is known about the normative microstructural organization of the human insular cortex and its relation to behavior. Precise quantitative *in vivo* characterization of the microstructural organization of the human insular cortex and links to its functional circuitry is critical for understanding its function and role in development and psychopathology.

The insular cortex is a structurally heterogenous brain region with a distinct cytoarchitectonic profile characterized by less differentiated cortical layers(1, 10, 11). Investigations of the microstructural features of the human insular cortex have been based solely on postmortem brains with small samples and limited behavioral characterizations. Insular cytoarchitectonic organization has been investigated using histological techniques over the last century(12–16). Stereological analyses of the insula have identified a cellular cortical architecture which differs considerably from the 6-layer granular architecture seen in most cortical areas(10, 17, 18). In a seminal study, Mesulam and Mufson(17) proposed the concept of “granularity”, based on the presence of an inner granular layer, as a key feature for identifying the anatomical subdivisions of the insular cortex. In the ensuing years, several histological studies(11, 12, 19) have focused on demarcating the microstructural properties of insular subdivisions but no consensus has yet emerged about their precise boundaries because of small sample sizes, limited insular divisions examined and high degree of variability across individuals. Thus far, investigations of the distinct structural features of the human insular cortex have been based solely on postmortem brains and have not been amenable to characterization using non-invasive brain imaging techniques. Quantitative modeling of non-invasive *in vivo* brain imaging are therefore critically needed to address this major gap.

Despite the lack of consistency and precision across previous histological studies, some general patterns have emerged regarding the general cytoarchitectonic organization of the insula. The ventral anterior insula has an agranular structure characterized by undifferentiated layers II/III and without a fully expressed layer IV compared to the fully developed granular cortex with a canonical 6-layer architecture. The dysgranular cortex has an intermediate profile that has been mainly observed in the dorsal anterior aspects of the insula(19). In contrast, large sections of the posterior insula show a canonical granular structure(19). A unique aspect of the neuronal organization of the human insula is the presence of von Economo neurons (VENs) that are present in the anterior aspects of the insula(9). VENs are projection neurons that differ from the typical pyramidal neurons by virtue of their large spindle shape and thick basal and apical dendrites(20, 21), which allow for speeded communication. The presence of VENs in the human insula is thought to contribute to its unique and important role in goal-directed behaviors and emotional regulation, through rapid processing of attentional, cognitive, interoceptive, emotional, and autonomic signals(4, 22, 23). The lack of tools for assessing morphological alterations in the insula *in vivo* and its relation to behavior has limited our understanding of the microstructural organization of the insula in health and disease.

The insular cortex is also functionally heterogeneous and integrates signals across its cognitive and affective subdivisions to support adaptive behavior. The anterior aspects of the insula are important for detection of salient external stimuli and for mediating goal-directed cognitive control while the posterior aspects are important for integrating autonomic and interoceptive signals(4). This functional organization is supported by a distinct pattern of long-range connections: the anterior insula is more strongly connected to brain areas important for cognitive control, most notably the dorsal anterior cingulate cortex (ACC) while the posterior insula has stronger links with subcortical and limbic regions important for emotion, including the amygdala and ventral striatum (2, 4, 24). Critically, the anterior insula and the dorsal ACC anchor the salience network (SN), a tightly coupled network that is among the most widely co-activated set of brain regions in all of human neuroimaging research(5–7). Remarkably, besides the insula, the only other brain region where VENs are known to be strongly expressed is the ACC(12, 25). The link between the microstructural organization of the insula and its functional circuit properties is currently not known. Furthermore, it is unclear whether these regional functional circuit properties are mirrored in the long-range connectivity of the insula in the context of the large-scale organization of AI-ACC circuits that anchor the SN. To the best of our knowledge there have been no histological investigations of the correspondence between cytoarchitecture features of the AI and ACC.

Given the challenges inherent in obtaining histological data from postmortem brains, the bulk of current research on the human insula has focused on its functional organization both in relation to regional activation by specific cognitive and affective tasks and its distinct patterns of functional connectivity with other brain regions(5, 26–28). Crucially, non-invasive voxel-wise quantitative mapping of functional brain connectivity has allowed researchers to identify distinct subdivisions within the insular cortex(26, 27, 29). Based on their unique fingerprints of connectivity, three distinct functionally subdivisions of the insular cortex have been identified: the dorsal anterior (dAI), ventral anterior (vAI), and posterior insula (PI) (9, 26). These subdivisions also show distinct patterns of intrinsic functional connectivity within the SN, with the dAI being more tightly linked to the ACC node of the SN(26). Whether functionally-defined regions of the insula and its functional circuits associated with the SN have distinct microstructural features is currently not known, and addressing this link has the potential to contribute to a deeper understanding of structure-function relations in the human brain.

Noninvasive *in-vivo* investigation of tissue microstructure is a key application of diffusion MRI (dMRI), but few studies have tapped its potential due to limitations of most previous dMRI techniques(30). A key discovery that now permits more precise microstructural analysis of human gray matter arose from the work by Latour et al(31) demonstrating the feasibility of analyzing cellular size in biological tissue with time-dependent diffusion MR. In subsequent studies, dMRI was used to determine sensitivity of dMRI signals in gray matter tissue(32, 33). More recently, dMRI has been used to characterize normal human brain maturation(34) and cortical microstructure in the preterm infants(35). Building on these studies, multi-shell models have recently been used to demonstrate regional variability in cortical microstructure(30, 36), as well as sensitivity of multi-shell dMRI for probing fine-scale and region-specific microstructure using high angular and spatial resolution diffusion MRI (37, 38).

Here we leverage recent advances in multi-shell dMRI acquisition protocols and recent signal reconstruction techniques(39, 40) to determine the microstructural features of insular cortex using the normalized Return to Origin Probability (RTOP) density index(41, 42). Crucially, as we demonstrate, RTOP is sensitive to cell size: the lower the RTOP, the larger the cell size. The first aspect of our study involved leveraging a large (N = 440) cohort of adult participants from the HCP. We evaluated RTOP across three major functional subdivisions of the human insula and demonstrate that they are microstructurally distinct with profiles consistent with its known agranular, dysgranular and granular organization. Based on observations that VENs have larger cell body than typical pyramidal neurons, and that VENs are more expressed in agranular and dysgranular, compared to granular, insula, we predicted differential levels of RTOP across insular subdivisions and smaller RTOP values in regions with agranular and dysgranular organization. We then examined whether the microstructural distinctions seen in the insular cortex are mirrored in the ACC node of the salience network, and demonstrate a strong link between functional circuits linking the insula and ACC and their microstructural organization. Finally, we examine whether variation in microstructural features of the insula are related to behavior and individual differences in cognitive control ability. Stability and cross-validation analysis are used to demonstrate the robustness of our findings.

The second aspect of our study focused on microstructural features of the macaque insular cortex, drawing on an extensive body of histological studies in non-human primates(15–17, 17, 43, 44). This allowed us to more precisely determine whether *in vivo* measurements using RTOP could reveal microstructural features consistent with the known agranular and dysgranular organization of the primate insula. Accordingly, we acquired data from two macaque monkeys using dMRI protocols similar to the HCP and deployed the same quantitative modeling approach as with the human dMRI data.

The third and final aspect of our study involved theoretical analysis and computer simulations based on known realistic 3-dimensional models of neuronal morphology to determine the sensitivity of RTOP measures to cell size(39, 41, 42), and in particular to the presence of large neurons such as the VENs. We synthesize findings from multilevel quantitative analysis and modeling to elucidate novel properties of the primate insular cortex and its relation to human cognitive abilities.

## Results

### Insula microstructure across its functionally defined subdivisions

Insula microstructure was characterized using RTOP(41) (see SI Materials for details of mathematical formulation). We focused on a widely used functional parcellation of the insula (26) consisting of the dorsal anterior insula (dAI), ventral anterior insula (vAI), and posterior insula (PI) in each hemisphere (**Figure 2A**). To determine whether RTOP values differed among the three insular subdivisions, we conducted an ANOVA with factors subdivision (vAI, dAI and PI) and hemisphere (left vs. right). We found a significant interaction between hemisphere and subdivision (F=157.13, *p*<5.0E-59) and significant main effects of subdivision (F=2850.3, *p*<4.7E-14) and hemisphere (F=60.76, *p*<4.7E-14). Post-hoc *t*-tests further revealed a gradient of RTOP values vAI < dAI < PI in both right and left hemisphere (all *ps*<2.0E-11, except *p*=0.036 in the case of right vAI < dAI), with vAI and PI having significantly smaller values in right, compared to the left, hemisphere (*ps*<1.2E-12) (**Figure 2B**).

**Figure 1.**
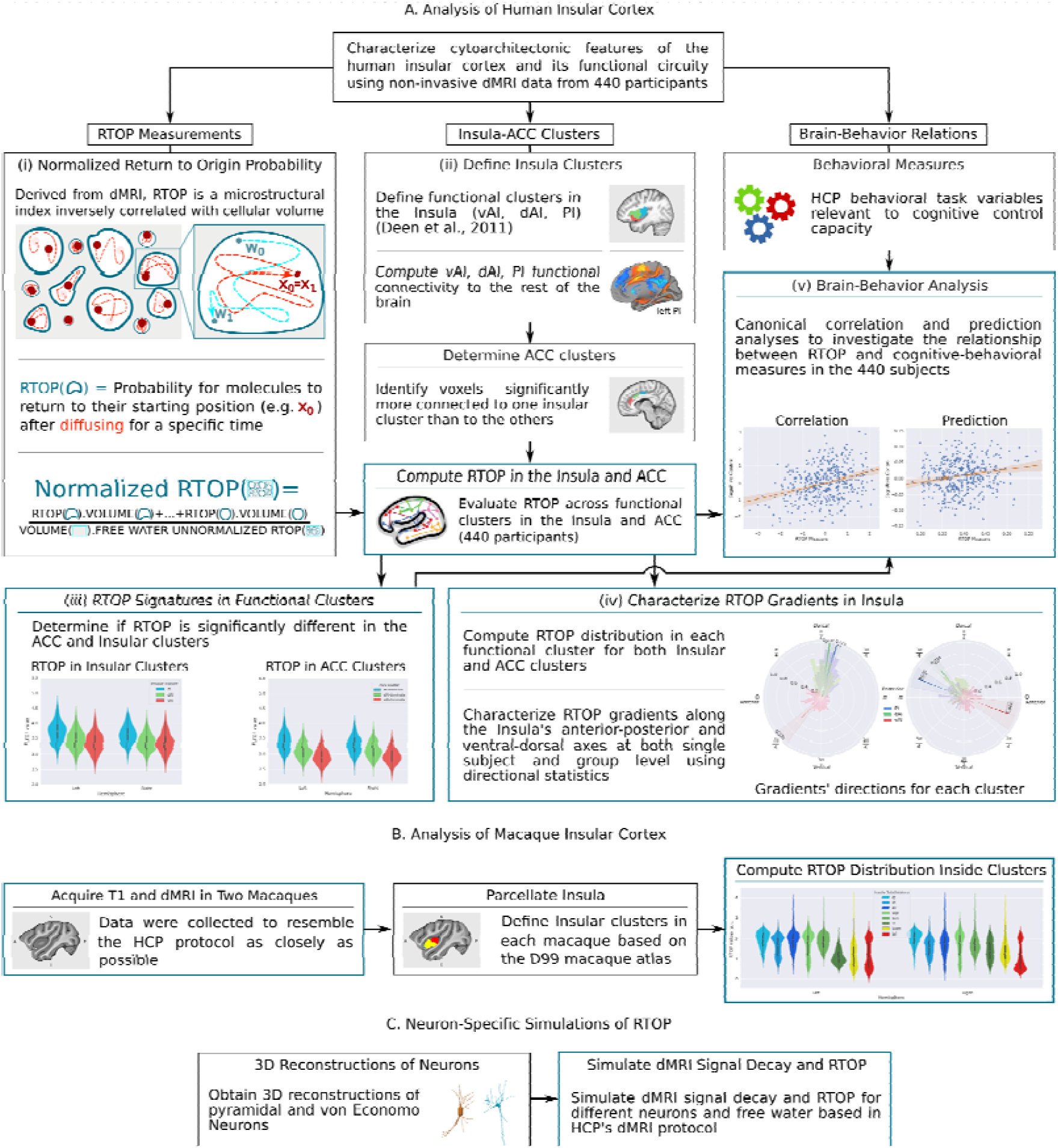
Flow chart illustrating data analysis pipeline. **(A)** Main components of human insula microstructure analysis using dMRI data from 440 Human Connectome Project (HCP) participants. Key steps include: **(i)** measurement of microstructure features based on Return to Origin Probability (RTOP), the ratio between the probability of molecules returning to their starting position in biological tissue versus free diffusion, **(ii)** demarcation of functional subdivisions in insula and its interconnected anterior cingulate cortex (ACC) subdivisions, which together anchor the salience network, **(iii)** computation of microstructure features of the insula within its functional subdivisions and its interconnected ACC subdivisions, **(iv)** computation of microstructural gradients along the anterior-posterior and dorsal-ventral axes of the insula, and **(v)** analysis of relation between insula microstructural organization and cognitive control abilities. **(B)** Main components of primate insula microstructure analysis using dMRI data from two macaques acquired with similar protocols used in the HCP. RTOP values were computed across granular and dysgranular subdivisions of the macaque insular cortex. **(C)** Computer simulations and theoretical analysis of RTOP illustrating sensitivity to cell size and presence of large neurons such as the VEN.

**Figure 2.**
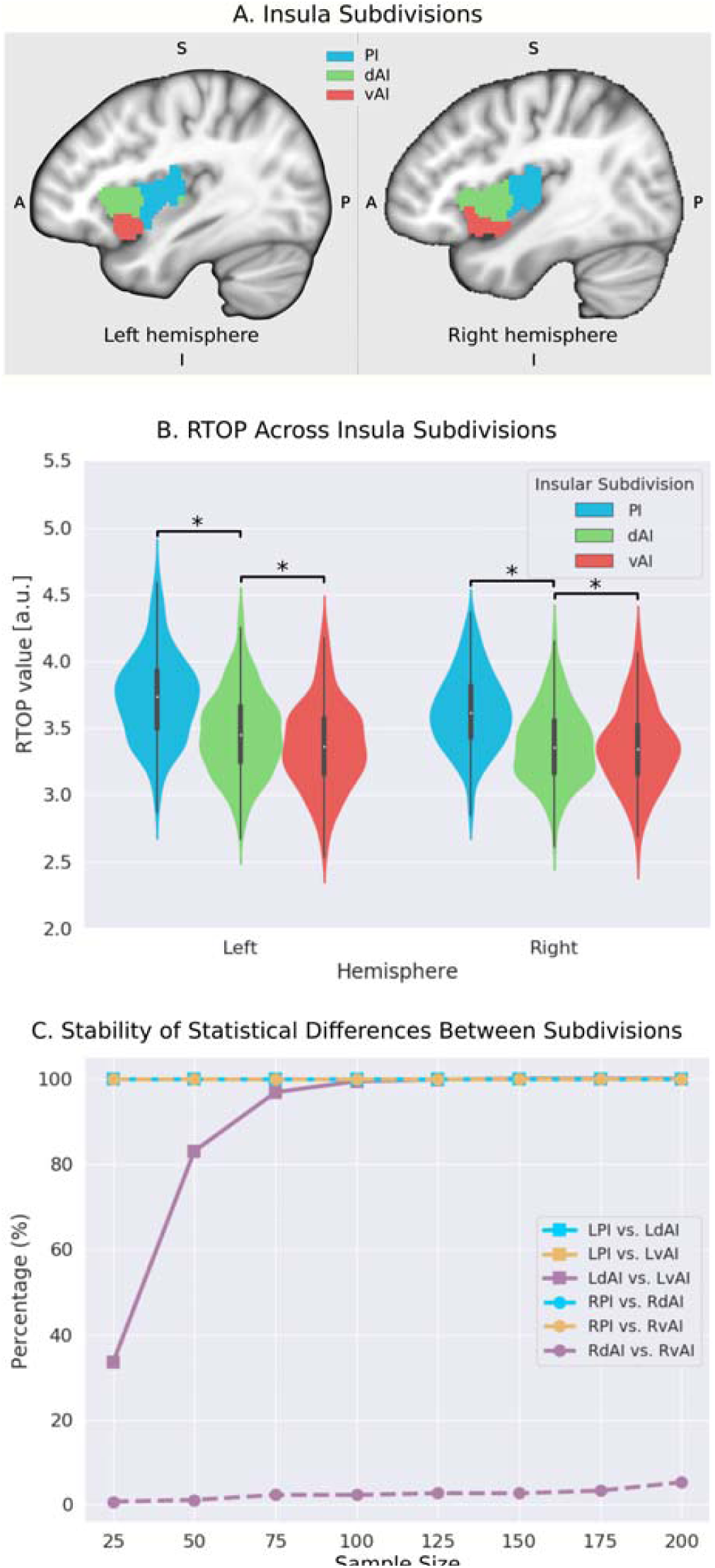
Microstructural properties of functional insular cortex subdivisions. **(A)** Functional subdivisions spanning the posterior insula (PI), dorsal anterior insula (dAI) and ventral anterior insula (vAI) (adapted from Deen et al). **(B)** RTOP is significantly different among PI, dAI and vAI in the left hemisphere and between PI and dAI or vAI in the right hemisphere (*p*<0.001, Bonferroni corrected). The right vAI has the smallest RTOP value among all the subdivisions (all *ps*<0.001, except *p*=0.036 for right vAI < dAI; Bonferroni corrected). **(C)** Stability of findings as a function of sample size. A sample size of N=25 was sufficient to achieve a stable differentiation (*p*<0.01) between PI and vAI in both hemispheres, while differentiating the vAI and dAI required a larger sample size. a PI: posterior insula; dAI: dorsal anterior insula; vAI: ventral anterior insula.

Next, we examined the stability of these findings as a function of sample size. We found that a sample size of N=25 was sufficient to achieve a stable differentiation (*p*<0.01) between PI and vAI in both hemispheres, while differentiating the vAI and dAI required a larger sample size of N=100 (**Figure 2C**).

Our findings provide robust evidence for distinct microstructural variations across these functional subdivisions of the insula, and suggests a close correspondence between the microstructural and functional organization of the insula.

### Gradient analysis of insula microstructural features along the anterior-posterior and dorsal-ventral axes

To further delineate the microstructural organization of the human insula, we conducted a detailed profile analysis and used directional statistics to characterize gradients in RTOP along its anterior-posterior and ventral-dorsal axes (**Figure 3**). **Figure 3A** shows isolevels of RTOP values in the insula at a group level. We found a low dispersion of RTOP gradients in both the left (0.18π ± 0.09 π, SEM=0.005) and right (0.23π ± 0.10 π, SEM=0.005) hemispheres. We then assessed the significance of this finding using the Rayleigh test, against the null hypothesis of a uniform directional distribution(45). We found strong evidence for both anterior-to-posterior and ventral-to-dorsal gradient in the left insula (ci=0.02π, Rayleigh statistic = 982, p < 1.0e-10, N=440, df=3) and a ventral-to-dorsal gradient in the right insula (ci = 0.03π, Rayleigh statistic = 628, *p* < 1.0e-10, N=440, df=3) (**Figure 3B-C, Table S1**). The confidence interval showed a low dispersion, indicating a consistent pattern of gradients across participants (**Table S1**). The lowest values of RTOP were localized to the vAI, convergent with findings from the analysis of the three functional subdivisions described above. Finally, analysis of microstructure isolines further revealed gradient ‘fingers’ from the vAI extending along a ventral-dorsal axis (**Figure 3D**). These results point to a consistent and reliable pattern of gradients and microstructural organization of the human insula along its anterior-posterior and dorsal-ventral axes.

**Figure 3.**
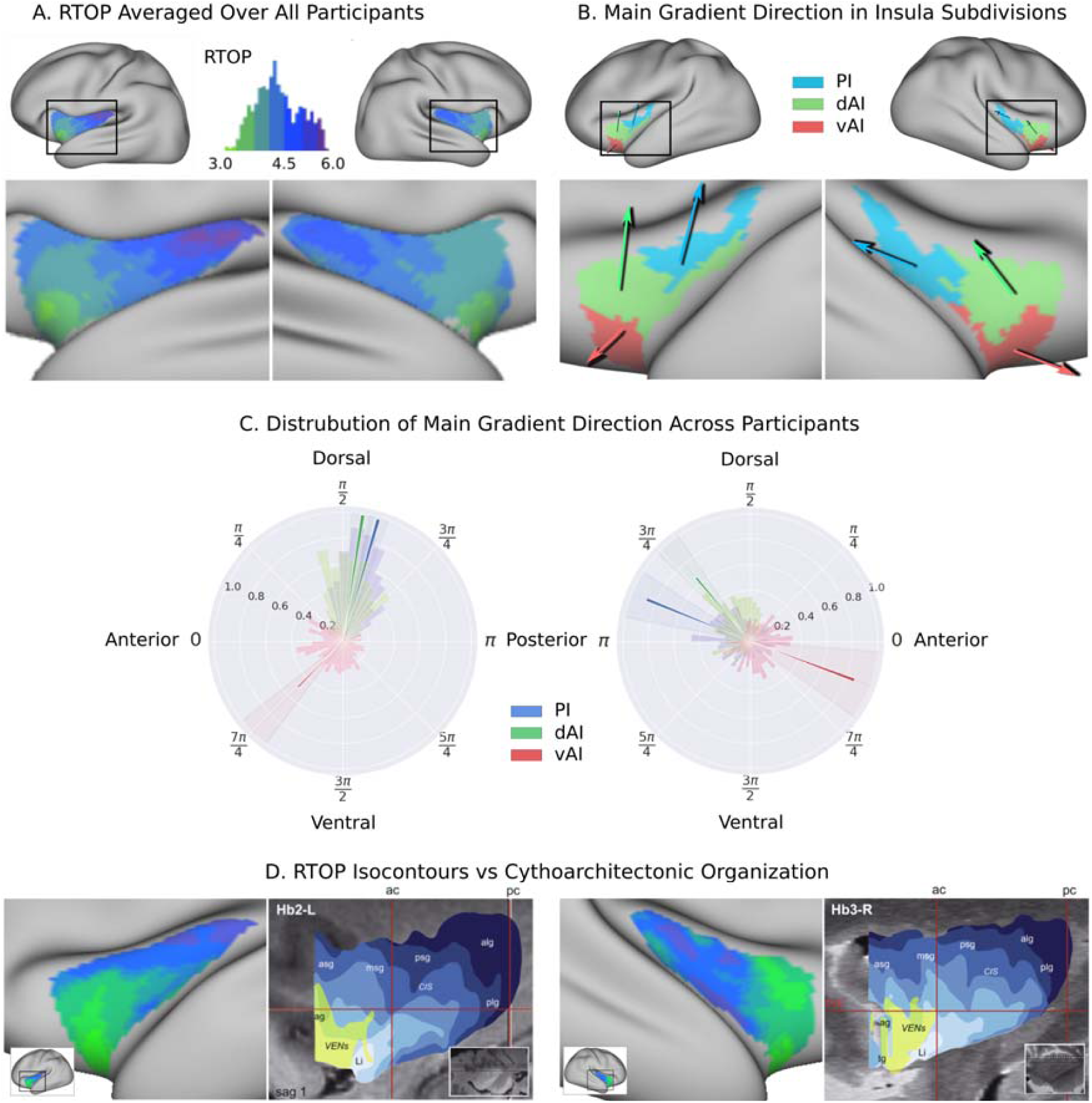
Insula microstructure gradients along its anterior-posterior and dorsal-ventral axes. **(A)** RTOP averaged over all participants (N=440) illustrates inhomogeneity in insula microstructure with a ventral anterior insula peak, and gradients along the anterior-posterior and dorsal-ventral gradients axes. Larger RTOP indicates smaller average compartments. There is a prominent gradient from the insular pole towards the posterior insular section. Note right hemisphere dominance. **(B)** Main gradient direction, computed using Rayleigh directional statistics in each functionally defined subdivision. The main directions show an anterior-to-posterior and inferior-to-superior RTOP organization in the left insular cortex and an anterior-to-posterior organization in the right insular cortex. The polar plots show the distribution of main gradient directions in each functional subdivision. **(C)** Gradient direction histograms. The mean direction is represented with solid lines on top of the distribution histogram; the shaded region represents the 95% confidence interval. For detailed statistics, please see **Table S1. (D)** Isocontours of the population-average RTOP (left) are closely aligned to cytoarchitectonic organization of the insula and VEN expression from studies of post-mortem brains (right, based on Morel et al. 2013).

### Microstructural features in the insula and ACC nodes of the salience network

The insular cortex and ACC are the two major cortical nodes of the salience network (SN)(2, 22). Individual functional subdivisions of the insula have preferential connections to different subdivisions of the ACC(26, 28, 46, 47), but it is not known whether they share similar microstructural features. To address this gap, we first conducted a seed-based whole-brain functional connectivity analysis, where seeds are the three functional subdivisions of the insular cortex in each hemisphere(26). Consistent with previous findings, the three insular subdivisions had distinct functional connectivity patterns with the ACC (**Figure 4A**). Specifically, the vAI showed stronger connection with the most anterior and ventral ACC (denoted as ACC-vAI), the dAI showed stronger connection with the middle and dorsal ACC (ACC-dAI), and the PI showed stronger connections with the posterior ACC (ACC-PI) (**Figure 4B**).

**Figure 4.**
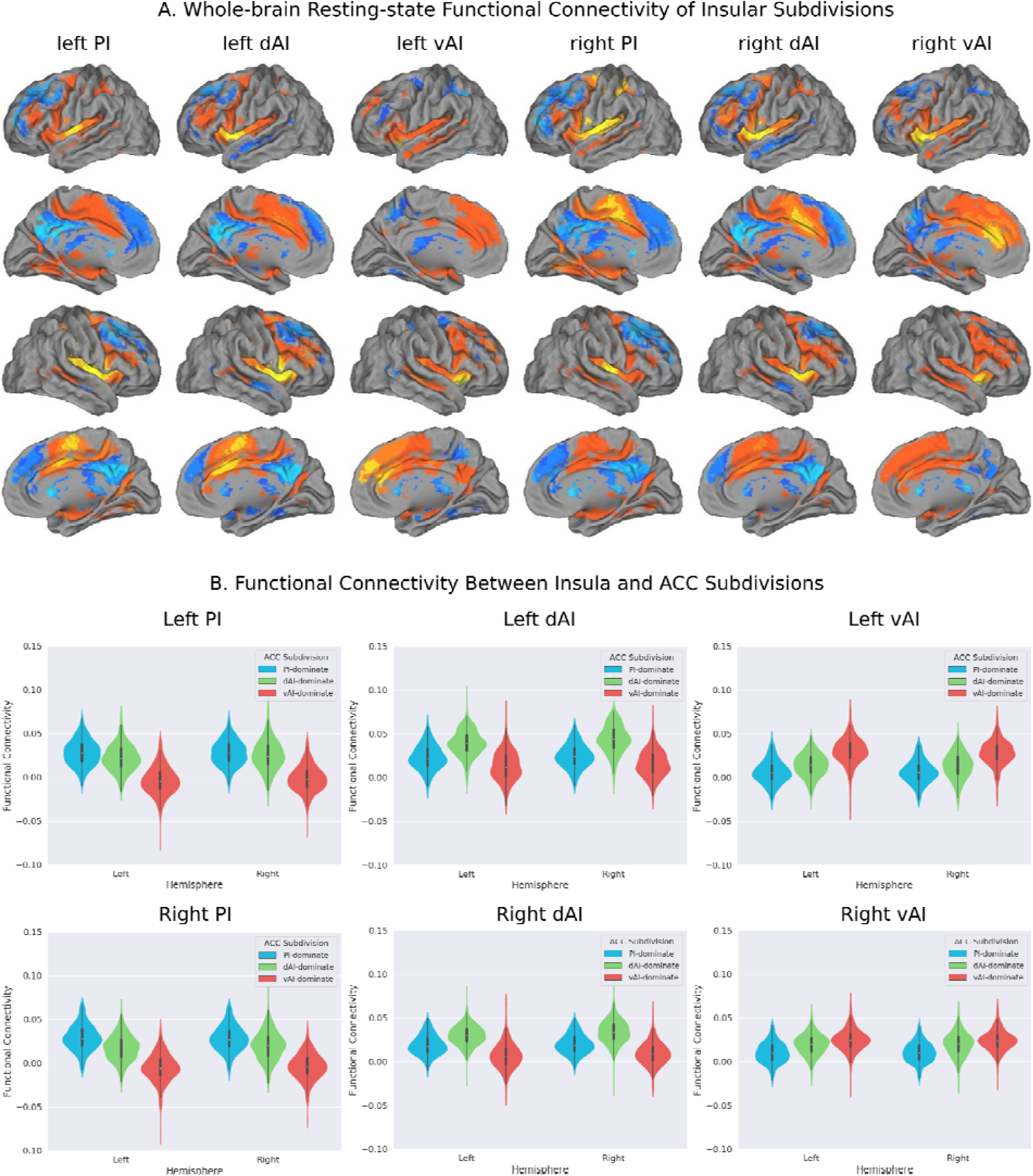
Intrinsic functional connectivity of insula subdivisions dAI, vAI and PI. **(A)** Whole-brain resting-state functional connectivity analysis revealed distinct functional connectivity patterns of the insula subdivisions, bilatera posterior insula (PI), dorsal anterior insula (dAI) and ventral anterior insula (vAI) (all *ps*<0.001, FDR corrected). Insular subdivisions were defined using an independent study (Deen et al., 2011). **(B)** RTOP differs across the ACC functional subdivisions linked to individual insular subdivisions.

Next, we created functional subdivisions within the ACC based on their differential connectivity patterns with the dAI, vAI and PI. First, we examined functional connectivity differences between each pair of seeds in each hemisphere (e.g. left PI > left dAI, thresholded (p<0.01, FDR corrected). We then applied a logical AND operation to identify voxels surviving the two binarized maps that indicated the voxel’s connectivity to one insula subdivision is significantly greater than the others (e.g. left PI > left dAI & left PI > left vAI). Finally, the resulting binarized maps were overlapped with an ACC mask (**Figure 5A**).

**Figure 5.**
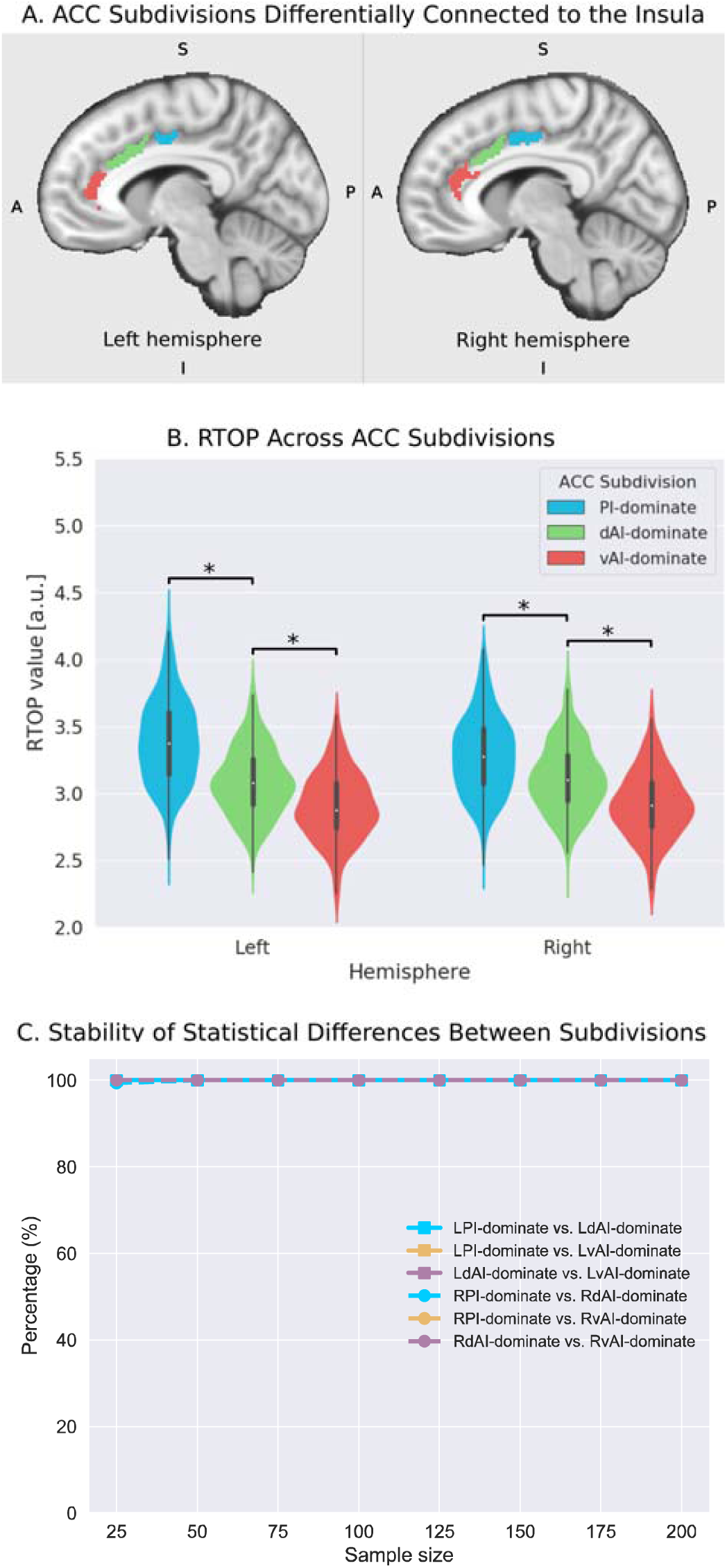
Microstructural properties of anterior cingulate cortex subdivisions which are differentially connected to functional subdivisions of the insula. **(A)** Illustration of ACC subdivisions. Each ACC subdivision preferentially connects to one of the three insular subdivisions defined in an independent study (Deen et al., 2011). ACC subdivisions showed significantly greater functional connectivity to one insula subdivision than the others, e.g (right PI > right dAI) & (right PI > right vAI) (all *ps*<0.01, FDR corrected). **(B)** RTOP values were significantly different among three ACC subdivisions in each hemisphere (*p*<0.001, Bonferroni corrected). The ACC subdivision differentially connected to vAI has smaller RTOP values than the other subdivisions (all *ps*<0.001, Bonferroni corrected). **(C)** RTOP differences among three ACC subdivisions were robust and reliable at sample sizes of N=25 or more. PI: posterior insula; dAI: dorsal anterior insula; vAI: ventral anterior insula.

To determine whether RTOP differs across the ACC functional subdivisions linked to individual insular divisions we conducted an ANOVA with factors subdivision (ACC-vAI, ACC-dAI and ACC-PI) and hemisphere (left vs. right). We found a significant interaction between subdivision (F=1106.9, *p*=2.93E-240) and hemisphere, and significant main effects of subdivision (F=1600.1, *p*=7.07E-293) and hemisphere (F=359.15, *p*=6.58E-59). Post-hoc paired *t*-tests revealed significant differences in RTOP: ACC-vAI < ACC-dAI < ACC-PI in both hemispheres (all *ps*<2.02E-16) (**Figure 5B**). Stability analysis demonstrated that these differences were highly reliable (*p*<0.01) for samples sizes > N=25. These results demonstrate that the microstructural organization of the ACC mirrors the microstructural organization of the three insula subdivisions with which it is differentially connected.

### Insula microstructure and relation to cognitive control ability

The human insula plays an important role in detection of salient external stimuli and in mediating goal-directed cognitive control(4). We investigated the relationship between microstructural properties of the insula and cognitive control ability, using canonical correlation analysis (CCA) with cross-validation and prediction analysis. Mean RTOP values from the six insular subdivisions, 3 in each hemisphere, were used to predict cognitive-behavioral measures associated with processing speed, working memory, response inhibition and cognitive flexibility. We found a significant relation between insula RTOP values and individuals cognitive control abilities (Pillai’s trace=0.21, *p*<0.01). The canonical weights of the 1^st^ latent variable in microstructural measures were significantly correlated with the canonical weights of the 1^st^ latent variable in behavioral measures (*r*=0.31, *p*<0.001, **Figure 6a; Table S2**). A leave-one-out cross-validation procedure further revealed that, based on microstructural properties of the insula, our CCA model could predict cognitive control ability on unseen data (*r*=0.18, *p*<0.001, **Figure 6b**), with the strongest predictive weights in the right AI.

**Figure 6.**
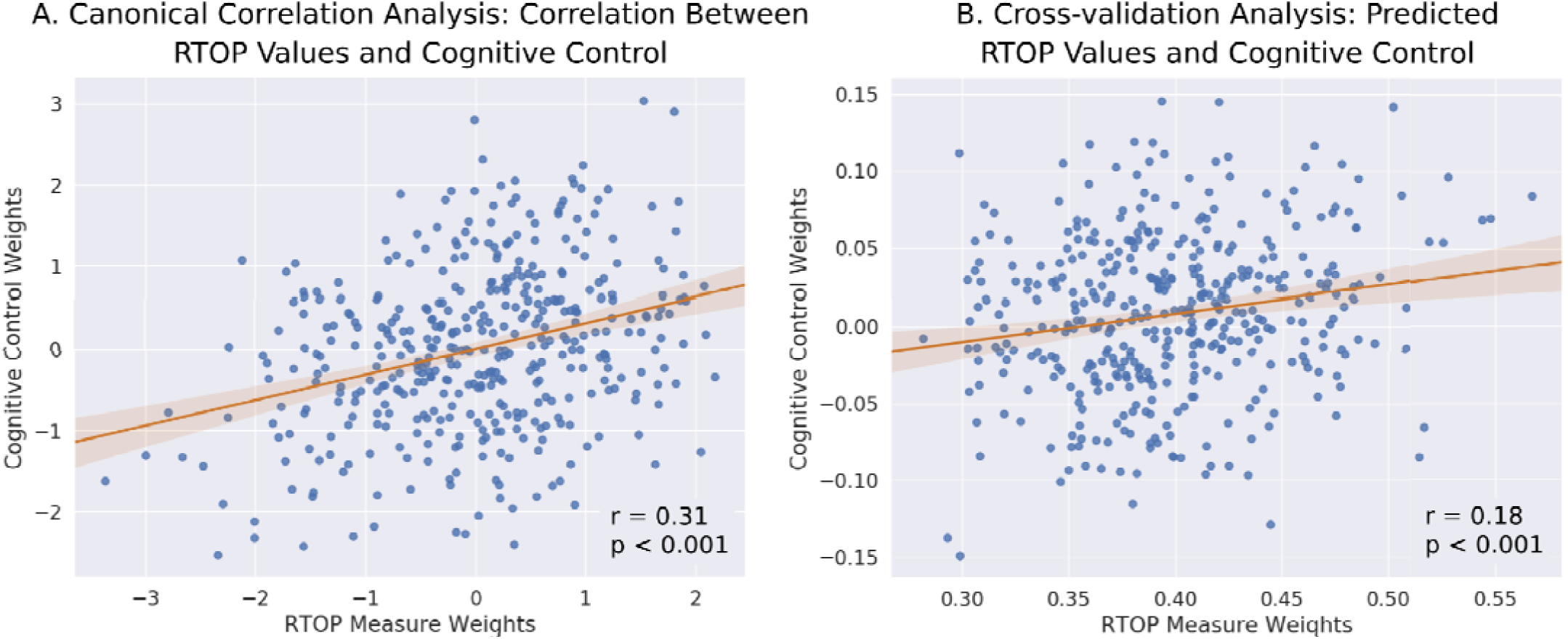
Insula microstructural features predict cognitive control abilities. **(A)** Canonical correlation analysis (CCA) revealed a relationship between RTOP and cognitive control measures. CCA weights of RTOP measures in Axis 1 has significant correlation with CCA weights of cognitive measures in CCA Axis 1. **(B)** Cross-validation analysis revealed that predicted CCA weights of RTOP and cognitive measures on unseen data are significantly correlated.

### Microstructural organization of insular cortex in macaque brain

Previous histological studies have demonstrated that the macaque insula can be clearly demarcated into agranular, dysgranular and granular subdivisions based on their unique cytoarchitectural properties with distinct profiles of VENs (43, 44) (**Figure 7A**). To investigate whether RTOP measures are sensitive to known cytoarchitectural features of the primate insula, we acquired dMRI data from two macaque monkeys, X77 and X181, using protocols similar to the HCP, and examined RTOP values in the known cytoarchitectonic subdivisions of the primate insula. We found that in both macaques, the lateral agranular insula (Ial) was the region with the lowest RTOP values. Specifically, in animal X77, the RTOP values inside the right Ial were significantly lower than other insular regions (**Figure 7B, Table S3**). In X77’s left hemisphere, the Ial had significantly lower RTOP than all other regions (two-sided t-test) except Agranular Insula (Ia) and the posterior-medial agranular insula (Iapm) (**Figure 7B, Table S3**). In X181, the left Ial showed significantly lower RTOP than other insular regions (**Figure 7B, Table S3**). Finally, the right Ial had lower RTOP than other regions, except for the Ia (**Figure 7B, Table S3**). Crucially, the granular insular cortex (Ig) contained the highest RTOP values in both monkeys. Ig also has significantly higher RTOP values than other insular regions, with the exception of the left intermediate agranular insula (Iai) in X77 (**Table S3**). These results demonstrate that the agranular insula shows the lowest RTOP, demonstrating (i) converging findings with known cytoarchitectonic studies of the insula in macaques and (ii) sensitivity of RTOP to average cellular size in the insula.

**Figure 7.**
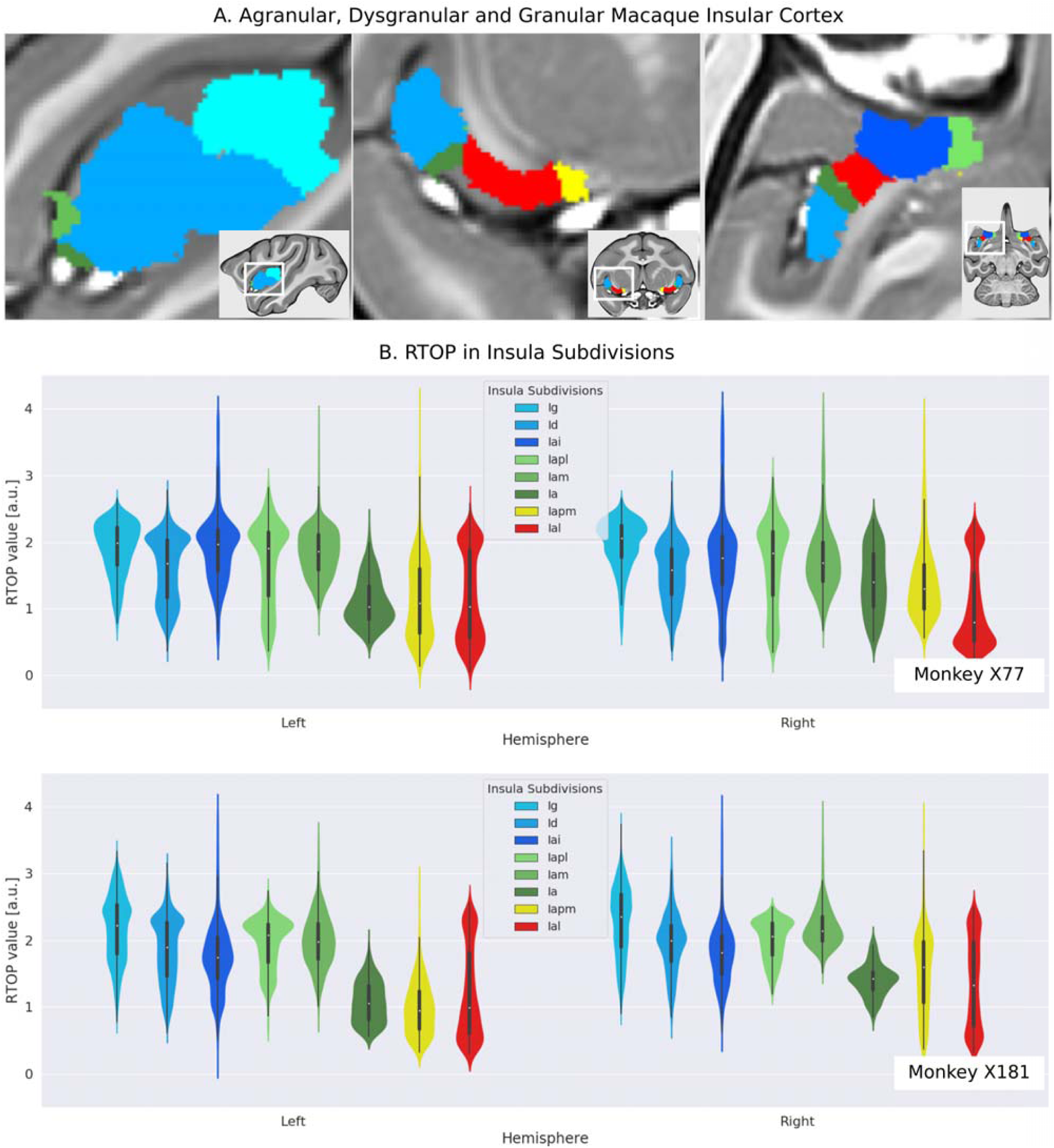
RTOP measurements validate the cythoarchitectonic organization of the macaque insular cortex *in vivo*. **(A)** Cytoarchitectonic subdivisions of the insula in the macaque monkey. **(B)** Distribution of RTOP values inside the cytoarchitectonic insular subdivisions for the two monkeys (X77 and X181). RTOP inversely correlates with expected average neuron size: RTOP is significantly lower in the lateral agranular insula (Ial) region (*p*<0.001) and adjoining posteromedial agranular insula (Iapm) and agranular insula (Ia), compared to other insular subdivisions. Granular insula (Ig) region has significantly higher RTOP values than other insular subdivisions, except for the intermediate agranular insula (Iai) in monkey X77 (*p*<0.001).

### Computational and 3D morphometric modeling of RTOP-derived microstructural features

We used computational modeling and theoretical analysis to examine RTOP sensitivity to cell size and the presence of VENs. First, analysis using simplified cell geometry revealed that RTOP scales as 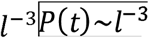, where *l* is the average characteristic length of the cellular compartment (see **Supplementary Materials**).

Second, we investigated whether RTOP can provide quantitative metrics of regional gray matter organization in multi-shell diffusion-weighted MRI (dMRI) HCP data. Briefly, we fit a regularized harmonic function representation(39, 40) to represent the dMRI signal. At each voxel, and for each participant we fixed model parameters using generalized cross-validation. We then calculated RTOP analytically from the underlying signal representation. In contrast to recent analyses of the multi-shell dMRI signal in the cortex(36), our model doesn’t require an inverse model fitting which is be prone to model degeneracies(48). We then tested the hypothesis that RTOP is sensitive to pyramidal neurons and VENs and that both induce an attenuation larger than the background signal to noise ratio. Based on simplified models of the pyramidal neuron and VEN somas as spheres(44) we found that the dMRI signal is sensitive to the presence of these neurons. Notably, enlarged VEN volume can induce signal decay of up to 88% in the HCP dMRI signals (see Supplementary Materials, **Figure S1**).

Next, to more precisely determine RTOP sensitivity to the presence of large neurons, we then used 3D digitally reconstructed models of fronto-insular human VEN and pyramidal neurons from NeuroMorpho.Org (12, 49), a Central Resource for Neuronal Morphologies. We simulated the HCP dMRI acquisition protocol (50) using a 3-D finite element model of the Block-Torrey equation (51). dMRI signals were estimated from 15 VENs and 9 pyramidal neurons. We found that, even at the lowest gradient strength used in the HCP protocol (50), 56 mT/m, i.e. b-value = 1,000 s/mm^2^, VENs induce a significant decay of 60% in the dMRI signal (**Figure 8**). Furthermore, VENs showed lower RTOP compared to pyramidal neurons (Mann-Whitney U test *U*=35, n_VEN_=15 n_pyramidal_=9, *p* < 0.05) (**Figure S2**).

**Figure 8.**
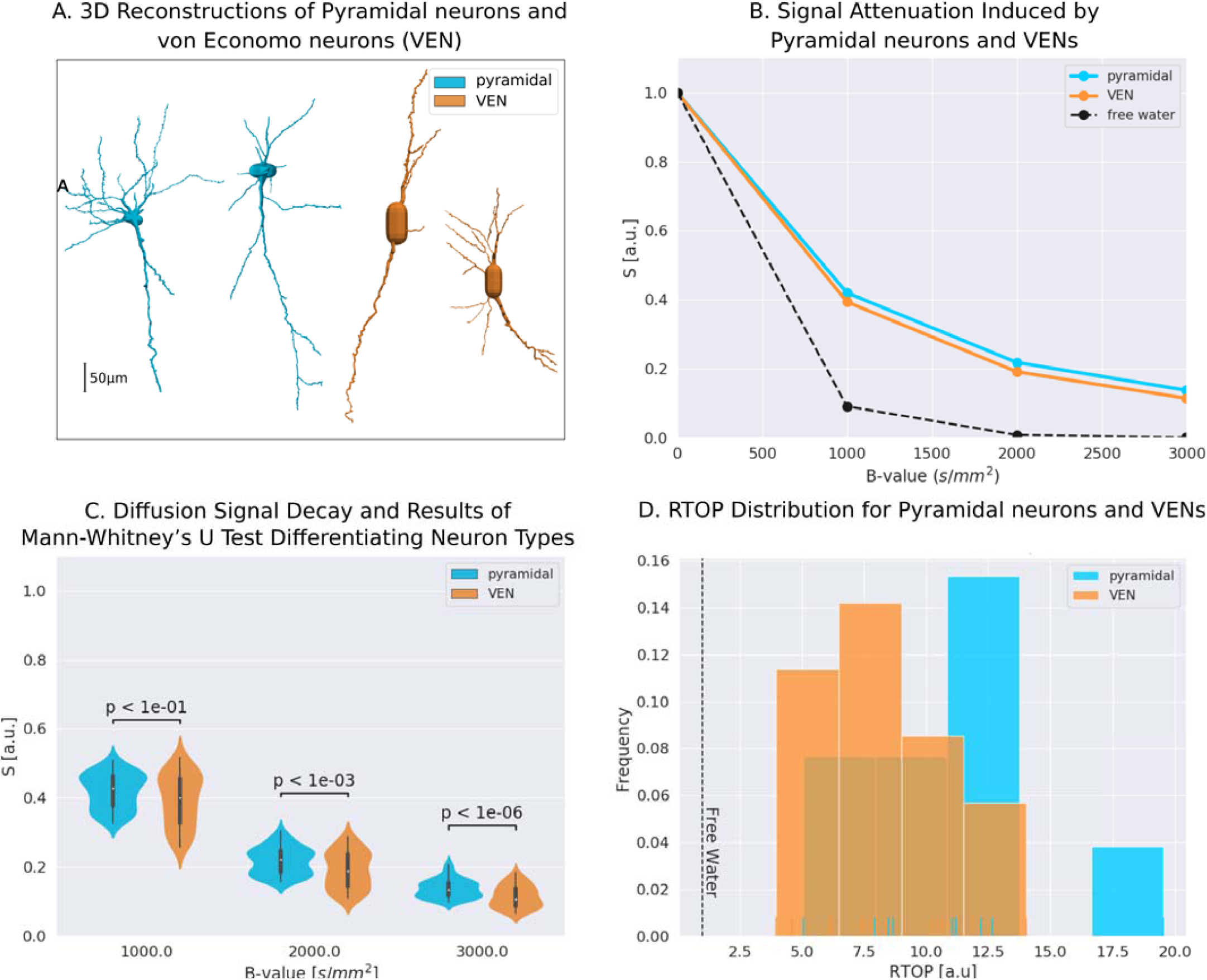
Dependence of RTOP on cell size and sensitivity to presence of VENs. **(A)** Illustration of three-dimensional reconstructions of VEN and pyramidal neurons from NeuroMorpho.Org used in the simulations. **(B)** Average dMRI signal attenuation as a function of acquisition parameters used in the human connectome project (HCP). Across all parameters, dMRI signal attenuation is larger in neuronal compartments compared to free water. **(C)** Signal attenuation is larger for VENs, compared to pyramidal neurons, leading to smaller RTOP values (*p* < 0.05). **(D)** Histogram of RTOP values for simulated VENs and pyramidal neurons.

Taken together, these results provide theoretical and *in silico* evidence that both pyramidal neurons and VENs attenuate the dMRI and modulate RTOP signals, and that RTOP measures are particularly sensitive to the presence of VENs.

## Discussion

We leveraged recent advances in diffusion weighted imaging of gray matter and large high-quality samples from the HCP to investigate the microstructural properties of the insular cortex and its macrofunctional circuits associated with the salience network. Novel quantitative modeling of *in vivo* dMRI and fMRI signals allowed us to probe links between the functional subdivisions of the insula and its microstructure properties, overcoming limitations of postmortem studies. We found that the human insular cortex is characterized by a systematic profile of microstructural variation across its distinct functional subdivisions. Beyond the boundaries of the functionally defined subdivisions, our analysis revealed gradients along the anterior-posterior axis as well as dorsal-ventral axis that are consistent with known cytoarchitectonic differences in the insula derived from studies of post-mortem brains(11, 52). Notably, we observed significant hemispheric differences with the right hemisphere showing lower RTOP values consistent with reports of larger neurons in the right hemisphere. Critically, the microstructural organization of the insula was mirrored in the dorsal ACC which together forms the backbone of the salience network, a system important for rapid orienting of attention and adaptive switching of cognitive control systems(53). Remarkably, microstructural properties of the insula were not only correlated with cognitive control abilities, but also predicted these abilities in a left-out sample. Our findings provide novel insights into human insular cortex architecture and its contribution to human cognition.

### Insula microstructure differs across functionally defined subdivisions

We used multi-shell dMRI data from a large cohort of 440 participants to characterize the microstructural properties of the human insular cortex in relation to its major functional subdivisions. There currently are no known landmarks that can distinguish different subdivisions of the insula on the basis of anatomical MRI data. To overcome this problem and provide a better link with its underlying functional organization, we took advantage of recent advances in functional parcellation of the human insular cortex(26, 27, 29). Resting-state fMRI studies have identified distinct functional subdivisions in the insular cortex based on differential patterns of intrinsic functional connectivity (26, 27, 29). These subdivisions have been widely used to determine the cognitive and affective role of the insula, and therefore serves as a basis for linking microstructure features of the insula with its functional subdivisions. The most widely used parcellation divides the insula into three functional clusters in each hemisphere: the dorsal anterior insula (dAI), ventral anterior insula (vAI), and posterior insula (PI) (**Figure 2**). Our analysis revealed robust evidence for distinct microstructural variations across these functional subdivisions of the insula. Specifically, in both the right and left hemispheres, the lowest RTOP values were in the vAI, followed by the dAI, and the largest in the PI. This pattern is consistent with the presence of VENs in the ventral and dorsal anterior insula, which form the agranular insular cortex in humans(9, 11, 25). Novel stability analysis revealed that this pattern was reproducible and reliable in samples of 50 or more. The stability of our findings based on functional subdivisions obtained from an independent study(26) thus suggest a close correspondence between functional organization of the insula and its microstructural organization.

### Anterior-posterior and dorsal-ventral gradients in insula microstructure

To further delineate the microstructural organization of the human insula, we conducted a detailed profile analysis to characterize gradients in RTOP along its anterior-posterior and ventral-dorsal axes. The lowest values of RTOP were localized to the vAI, convergent with findings from the analysis of the three functional subdivisions described above. We found strong evidence of an anterior-to-posterior and ventral-to-dorsal gradient in each subdivision of both the left insula and the right insula (**Figure 3B-C**). Analysis of microstructure isolines revealed gradient ‘fingers’ from the vAI extending along a ventral-dorsal axis. This pattern is strikingly consistent with findings from known cytoarchitectonic features in the insula derived from studies of post-mortem brains(11). The vAI regions where we observed the largest cell size (lower RTOP) is also consistent with histological studies that have found the highest density of VENs in this region(11, 12, 44) (**Figure 3D**).

The anterior-posterior-ventral-dorsal gradients and the convergent findings from the three distinct functional subdivisions provides new quantitative insights into the microstructural organization of the human insular cortex. The precise anatomical boundaries of the agranular, dysgranular, and granular architectonic areas within the human insula are not known, and differences in stereological methods and criteria have led to different segmentation schemes of the insular cortex in primates(54, 55). However, there is general consensus that the anterior and most of the mid insula areas are agranular or dysgranular while the posterior most aspects are granular(9). The spatial gradient of RTOP in the insular cortex along the dorsocaudal-rostroventral axis provides new metrics for noninvasive *in vivo* analysis of the general cytoarchitectonic organization in the insular cortex in the human brain and is an advance over previous studies that have thus far been based on invasive histological and electrophysiological data of human and non-human primates(11, 13, 15, 17, 56).

### Hemispheric asymmetry in microstructural organization of the human insula

Our analysis of the left and right insula revealed a prominent hemispheric asymmetry in microstructure. RTOP values were significantly lower in the right hemisphere, with the lowest values among all six subdivisions (three in each hemisphere) being localized to the vAI in the right hemisphere. This is consistent with reports of larger VEN in the right hemisphere in human postmortem data(12). Similarly, in the macaque insula, a comparison of the number of VENs in the left and right AI revealed a significantly higher number of VENs in the right anterior AI (44), in accordance with the asymmetry reported in the human brain(12). Interestingly, this asymmetry parallels the differential functional role and engagement of the right AI in cognitive and emotion control tasks such as the Go-Nogo, Stop signal and Emotional Stroop tasks in the human brain(5, 57, 58). Consistent with these observations, the right AI has been shown to exert significant causal influences on multiple other brain areas in a wide range of cognitive control tasks(5, 57– 59) and lesions to this region are known to impair cognitive control(60). Hemispheric asymmetry in microstructural organization of the insula, and its putative links with differential expression of VENs are consistent with reports of a left-right functional asymmetry in the insula(61). Specifically, it has been suggested that homeostatic afferent, including hot and cold pain, muscle and visceral pain, sensual touch and sexual arousal all produce strong right-lateralized activation in the right AI(62). Heartbeat-related evoked potentials and interoceptive awareness of heartbeat timing, arising from ‘sympathetic’ homeostatic afferent activity are also associated with AI activity(23). We suggest that hemispheric asymmetry in microstructural organization of the insula may contribute to lateralization of function and in particular the differential role of the right insula in monitoring internal bodily states and subjective awareness across a wide range of cognitive and affective processing tasks(62).

### Linked microstructural features in the insula and ACC nodes of the salience

The insular cortex together with the ACC are the two major cortical nodes that anchor the salience network (SN)(2, 22). Noninvasive brain imaging studies using both fMRI and DTI have shown that the insula and ACC are strongly connected functionally and structurally(5, 12, 22, 53, 63–65), together forming the backbone of the SN. Previous studies have shown that individual functional subdivisions of the insula have preferential connections to different subdivisions of the ACC(26, 28, 46, 47). However, it is not known whether functional subdivisions within the ACC share the similar microstructural properties to the insular subdivisions they connect to. In a significant advance over previous research, we found a close correspondence between microstructural organization of the insula and its interconnected ACC subdivisions. First, using whole-brain functional connectivity analysis we discovered three dorsal ACC subdivisions each with a pattern of preferential functional connectivity with the three insula subdivisions - vAI, dAI and PI. Crucially, as with the three insula subdivisions, we found that these three functionally-defined dorsal ACC subdivisions themselves have distinct microstructural organization. Moreover, the underlying pattern is characterized by a one to one mapping between their respective RTOP values. The ACC subdivision with low RTOP was strongly connected to the insular subdivision with low RTOP, and the ACC subdivision with high RTOP is strongly connected to the insular subdivision with high RTOP. Our findings suggest a strong link between functional circuits linking the insula and ACC and their microstructural organization.

Our findings provide novel insights into the link between the functional organization of the SN and its microstructural features. Although there have been no histological investigations of the correspondence between cytoarchitecture features of the AI and ACC, it is known that both brain regions are unique in their expression of VENs(12, 25, 66). Fronto-insular VENs in humans are known to project to the dorsal ACC(4, 12), and tract-tracing studies in macaques have shown that the ventral anterior insula receives input from pyramidal neurons in ipsilateral dorsal ACC(17). Our findings help link microstructural features of the insula with its macroscopic functional connectivity with the ACC for the first time. The correspondence between microstructural and large-scale functional connectivity profiles across the three insula and ACC subdivisions suggests that local and large-scale circuit features may together contribute to the integrity of the SN.

### Insula microstructure predicts cognitive control ability

The human insula has been implicated in a broad range of cognitive functions(2, 3, 5, 20, 67), but links between its microstructural features and individual differences in cognitive control abilities have not been previously examined. Our analysis revealed an association between the microstructural properties of the insula and individual differences in cognitive control ability. Remarkably, machine learning algorithms and cross-validation leveraging the large HCP sample size also revealed that cognitive control abilities could be predicted in a left-out sample with the strongest predictive weights in the right anterior insula. Our findings provide novel in vivo evidence that the microstructural integrity of the insula is crucial for implementing cognitive control. Gray matter lesions and insults to white matter pathways associated with the insular cortex and pathways linking it to the salience network have also been shown to impair cognitive control ability(60, 68, 69). In particular, the AI, the key node in the salience network, is one of the most consistently activated brain regions during tasks involving cognitive control(5, 6). Moreover, the strength of functional and anatomical connectivity between anterior insula and other cognitive control regions is modulated by cognitive demands(57–59, 67, 70–72). Aberrant connectivity of the anterior insula has been also associated with cognitive deficits in psychiatric disorders(73–75). Integrating these and other related findings, a prominent neurocognitive model has proposed that the salience network, especially the anterior insula, plays an important role in dynamically switching between other core brain networks to facilitate access to cognitive resources(2). A unique feature of the human insular cortex that is thought support its role in fast switching is the presence of VENs, whose large axons could be fundamental neuronal basis for rapid signal relay between AI and ACC as well as other brain networks. Our findings provide convergent support for this hypothesis and demonstrate a link between the unique microstructural features of the anterior insula and cognitive control function in humans, and they provide new quantitative metrics for investigating multiple psychiatric and neurological disorders known to impact the insula transdiagnostically(8).

### RTOP captures known microstructural features of the agranular insular cortex in macaques

Because of the paucity of histological data from the human insular cortex, we then sought to determine whether *in vivo* measurements using RTOP can reveal known microstructural features of the macaque brain. Recently, Evrard and colleagues have provided a detailed account of the macaque insula architecture which can be summarized as follows: agranular areas contain VENs, but no granule cells while granular insula regions have a large number of granule cells but few, if any, VENs(43). Histological studies in the macaque brain have shown that VENs are on average 50% and 70% larger than local pyramidal neurons and are concentrated within lateral agranular insula(43) (Ial). Consequently, we predicted that compared to granular insular cortex, RTOP values would be lower in agranular insula, reflecting the presence of large neurons such as the VENs. We found that RTOP values are indeed significantly lowest in the Ial in both right and left hemispheres, with more prominent effects in the right hemisphere where the right Ial had lowest values compared to the rest of the right insular cortex. In the left insular cortex, the lowest values of RTOP were found in the Ial, agranular insula (Ia) and posterior-medial agranular insula (Iapm), the three agranular regions (Ia, Iapm, Ial) which adjoin each other (**Figure 7**). These findings are consistent with histological findings that the agranular cortex contains a greater population of VEN cells and a smaller population of small granule cells(43). The presence of larger VENs increases the average cell size, thus decreasing restriction on the diffusion of water molecules, thereby resulting in lower RTOP values. Taken together, our analysis of the macaque brain provides new evidence that RTOP is sensitive to the presence of VENs and, critically, that it reflects the known cytoarchitectonic organization of the primate insular cortex.

### Computational and morphometric modeling reveals that microstructural measures are sensitive to large neurons and VENs

Finally, we used computational modeling and theoretical analysis to demonstrate RTOP sensitivity to cell size and the presence of VENs. In biological tissue, the dMRI signal(31) and RTOP density are known to be sensitive to volume, surface-to-volume ratio(41, 42), and mesoscopic organization of cells(76). Three different analyses yielded consistent findings linking RTOP with cell size. First, theoretical analysis using simplified cell geometry revealed that RTOP scales as 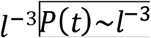, where l is the average characteristic length of the cellular compartment. Second, based on simplified models of the pyramidal neuron and VEN somas as spheres(44) we found that the dMRI signal is sensitive to the presence of these neurons with reductions in RTOP of up to 88%. Third, using realistic models of fronto-insular human VEN and pyramidal neurons from NeuroMorpho.Org, we found that VENs induce a significant decay of 60% in the dMRI signal. Furthermore, using extensive simulations we verified that compared to soma, the neuronal cell body, dendrites and axons, have a minimal effect on RTOP signals, consistent with previous reports(77). These quantitative results demonstrate that quantitative analysis of multi-shell dMRI data provides reliable and replicable noninvasive *in vivo* measures of gray matter microstructure and sensitivity to the presence of large neurons such as the VENs.

### Conclusion

Our novel quantitative analysis of multi-shell dMRI data provides reliable and replicable noninvasive *in vivo* measures of gray matter microstructure. We identified several unique microstructural features of the human insula and functional circuits associated with the salience network. Convergent findings from two macaques revealed microstructural features consistent with the known granular and dysgranular cytoarchitectural features of the primate insular cortex. Together with computational modeling of dMRI signals, results point to a gradient profile consistent with the presence of large VENs in the agranular anterior insula and histological data. Crucially, microstructural properties of the insular cortex are behaviorally relevant as they predicted human cognitive control abilities, in agreement with its crucial role in adaptive human behaviors. Our study provides a novel template for non-invasive investigations of microstructural heterogeneity of the human insula and the salience network, and how its distinct organization may impact human cognition, emotion and interoception. Our findings also open new possibilities for probing psychiatric and neurological disorders that are known to be impacted by insular and cingulate cortex dysfunction, including autism, schizophrenia, depression and fronto-temporal dementia (56, 78–81).

## Methods and Materials

### Human and monkey datasets

Data acquisition for the Human Connectome Project (HCP) was approved by the Institution Review Board of the Washington University in St. Louis (IRB #201204036), and all open access data were deidentified. We also used data from two male rhesus macaques (Macaca mulatta). The Princeton University Animal Care and Use Committee approved all procedures, which conformed to the National Institutes of Health guidelines for the humane care and use of laboratory animals. Details of the data acquisition, preprocessing and analysis steps are in the Supplementary Materials, see also **Figure 1**.

### HCP diffusion MRI data acquisition and preprocessing

Minimal preprocessed diffusion MRI data(50) from 440 participants (22-36 years old, 261 female/179 male) were obtained from the HCP Q1-Q6 Data Release. Details of the acquisition protocol and preprocessing steps are described by Sotiropoulos et al(50).

### Computing RTOP from diffusion MRI data

The dMRI signal characterizes the probability that water molecules within a voxel experience a net displacement after a given diffusion time. This enables the measurement of the probability that, after a diffusion time t, water in a voxel are in the vicinity of their starting position, namely the RTOP density(41). To compute RTOP we evaluated a regularized representation of the signal based on the Mean Average Propagator formalism with Laplacian regularization(40) (MAPL) included in the dipy open-source software package (http://nipy.org/dipy). The regularization parameter was selected through generalized cross validation and RTOP was computed analytically from the fitted MAPL parameters. Finally, to render the Return-To-Origin probability comparable across subjects, we normalized it by the average ventricular Return-To-Origin probability of each subject’s cortico-spinal fluid in the ventricles. In this way the normalized RTOP quantifies the enhancement ratio over free diffusion(41, 42) specific to each subject.

### RTOP stability analysis

To evaluate stability of our findings regarding RTOP differences between ROIs, we used subsampling procedures and determined minimum sample sizes that consistently reproduced findings.

### RTOP gradient analysis

To determine whether RTOP values followed an anterior-to-posterior and inferior-to-superior cytoarchitectonic organization of the insular cortex(1, 9, 11), we computed a RTOP gradient field along the insular surface of each participant. To assess that the main gradient directions obtained for each participant were following a common direction, their mean and its confidence interval was computed. Furthermore, we used a Rayleigh test to reject the hypothesis that participants’ main gradient directions were uniformity distributed.

### HCP resting-state fMRI data acquisition and processing

Minimal preprocessed resting-state fMRI data (session: rfMRI_REST1_LR) from 440 participants was obtained from the HCP Q1-Q6 Data Release. During scanning, each participant had they eyes fixated on a projected crosshair on the screen. Spatial smoothing with a Gaussian kernel of 6mm FWHM was first applied to the minimal preprocessed data to improve signal-to-noise ratio as well as anatomy correspondence between individuals. A bandpass temporal filtering (0.008 Hz < f < 0.1 Hz) was then applied.

### fMRI connectivity analysis

Seed-based functional connectivity analyses were conducted to examine whole-brain connectivity pattern of each insular subdivision. First, time series across all the voxels within each insular subdivision were extracted and averaged. The resulting averaged time series were then used a covariate of interest in a linear regression of the whole-brain analysis. A global time series, computed across all brain voxels, along with six motion parameters were used as additional covariates to remove confounding effects of physiological noise and participant movement. Linear regression was conducted at the individual level. A group map was generated using one-sample *t* tests (*p*<0.01, FDR corrected). Last paired *t* tests were applied at the group level to identify brain regions significantly more correlated with one subdivision than the other (*p*<0.01, FDR corrected).

### Functional subdivisions in anterior cingulate cortex

fMRI connectivity analysis was used to identity functional subdivisions of the ACC that were differentially connected to each of the insular subdivisions. First, maps of paired comparison between functional connectivity of different insular subdivisions were thresholded (*p*<0.01, FDR corrected) and binarized. Second, logical AND operation was applied to identify voxels surviving two paired comparisons for an insular subdivision versus the others (e.g. left PI > left dAI & left PI > left vAI). Last, the resulting mask was overlapped with an ACC mask from Harvard-Oxford Probabilistic Atlas of Human Cortical Area (http://www.cma.mgh.harvard.edu/fsl_atlas.html).

### Relation between insula microstructure and behavior

The relationship between brain and behavioral measures was examined using Canonical Correlation Analysis (CCA) and a cross-validation with prediction approach(82). Brain measures consisted of RTOP values for each subdivision of the insular cortex in each hemisphere (6 variables in total). Behavioral measures consisted of in- and out-scanner variables that are highly relevant to cognitive control capacity. In-scanner behavioral task measures included n-back working memory task accuracy and reaction time (RT), relational task accuracy and RT, and gambling task percentage accuracy and RT for larger choice. Out-of-scanner behavioral measures consisted of performance on List sorting, Flanker, Card sorting, Picture sequence tasks from the NIH toolbox as well as the processing speed. Together, there were 11 behavioral measures. Prediction analysis was performed using leave-one-out cross-validation. Pearson’s correlation was used to the evaluate the correlation between the predicted brain and behavioral measures.

### Macaque MRI data acquisition and processing

We scanned two male rhesus macaques (Macaca mulatta, ages = 4 yrs, body weights = 5.6/6.8 kg). For all scan sessions, animals were first sedated with ketamine (10mg/kg IM) and maintained with isoflurane gas (2.5-3.0%) using an MR-compatible anesthesia workstation (Integra SP II, DRE Inc, Louisville KY). Data were collected to resemble the HCP dMRI protocol as close as possible. Specifically, we acquired the whole brain on a Siemens 3T MAGNETOM Prisma (80 mT/m @ 200 T/m/s gradient strength) using a surface coil (11cm Loop Coil; Siemens AG, Erlangen Germany) secured above the head. Nine T1-weighted volumes were collected for averaging to obtain a high-quality structural volume and diffusion images were acquired using a double spin-echo EPI readout pulse sequence. Four datasets of 271 gradient directions were collected using a monopolar gradient diffusion scheme with b values distributed optimally across 3 spherical shells. All macaque MRI data will be/is publicly available on the NKI PRIMatE Data Exchange database(83). The T1-weighted images were registered to the D99 atlas through the NMT—NIMH Macaque Template allowing to subdivide the insula into granular, dysgranular, and agranular areas along with smaller subdivisions of each parcel. RTOP was computed using similar procedures as the human dMRI data.

### Computational modeling and simulation of dMRI signals using 3-dimensional microscopic reconstructions of VEN and pyramidal neurons

We performed simulations on 3-dimensional microscopic reconstructions of VEN and pyramidal neurons(12). Our simulation followed the same acquisition parameters as the HCP dMRI protocol. Specifically, finite-element-based simulations were performed using an open-source package for computational diffusion MRI developed in the FEniCS(85). The methods and software are described by Nguyen et al(51). Simulations were conducted for the whole neurons as well as for the soma and proceedings separately. Finally, RTOP was computed using similar procedures as the human dMRI data. All simulation data will be made available upon publication.

## Funding

### Author contributions

Conceptualization, Supervision, Project Administration, and Funding Acquisition: V. M and D. W; Methodology: D. W., V. M., G. G., J. L., W. C.; Investigation: V. M., W. C., G. G., D. W.; Resources and Data Curation: M. P., G. G., W. C., D. W.; Visualization: G. G.; Formal Analysis: D. W., V. M., G. G., J. L., W. C; Writing: V. M., G. G., W. C., D. W.

### Data and materials availability

All data is available in public databases. The human data comes from the Human Connectome Project, the primate data is available at the INDI Primate Data Exchange, and the three-dimensional neuronal models are available from the NeuroMorpho website.

### Code Availability

All custom code will be available on GitHub upon manuscript publication and will have a DOI-providing submission in Zenodo. All code were developed based on open-source, publicly available software packages.

## References

1. Nieuwenhuys R (2012) The insular cortex: a review. Prog Brain Res 195:123–163.

2. Menon V, Uddin LQ (2010) Saliency, switching, attention and control: a network model of insula function. Brain Struct Funct 214(5–6):655–667.

3. Uddin LQ (2015) Salience processing and insular cortical function and dysfunction. Nat Rev Neurosci 16(1):55–61.

4. Craig ADB (2009) How do you feel--now? The anterior insula and human awareness. Nat Rev Neurosci 10(1):59–70.

5. Cai W, Ryali S, Chen T, Li C-SR, Menon V (2014) Dissociable roles of right inferior frontal cortex and anterior insula in inhibitory control: evidence from intrinsic and task-related functional parcellation, connectivity, and response profile analyses across multiple datasets. J Neurosci Off J Soc Neurosci 34(44):14652–14667.

6. Swick D, Ashley V, Turken U (2011) Are the neural correlates of stopping and not going identical? Quantitative meta-analysis of two response inhibition tasks. NeuroImage 56(3):1655–1665.

7. Levy BJ, Wagner AD (2011) Cognitive control and right ventrolateral prefrontal cortex: reflexive reorienting, motor inhibition, and action updating. Ann N Y Acad Sci 1224:40–62.

8. Goodkind M, et al. (2015) Identification of a Common Neurobiological Substrate for Mental Illness. JAMA Psychiatry 72(4):305.

9. Namkung H, Kim S-H, Sawa A (2017) The Insula: An Underestimated Brain Area in Clinical Neuroscience, Psychiatry, and Neurology. Trends Neurosci 40(4):200–207.

10. Mesulam MM, Mufson EJ (1982) Insula of the old world monkey. I. Architectonics in the insulo-orbito-temporal component of the paralimbic brain. J Comp Neurol 212(1):1–22.

11. Morel A, Gallay MN, Baechler A, Wyss M, Gallay DS (2013) The human insula: Architectonic organization and postmortem MRI registration. Neuroscience 236(C):117–135.

12. Allman JM, et al. (2010) The von Economo neurons in frontoinsular and anterior cingulate cortex in great apes and humans. Brain Struct Funct 214:495–517.

13. Brodmann K (1909) Vergleichende Lokalisationslehre der Grosshirnrinde in ihren Prinzipien dargestellt auf Grund des Zellenbaues (Springer).

14. Von Economo C, Koskinas GN, Triarhou LC (2008) Atlas of cytoarchitectonics of the adult human cerebral cortex (Karger, Basil⍰ New York). 1st English ed.

15. Mesulam M-M, Mufson EJ (1985) The Insula of Reil in Man and Monkey. Association and Auditory Cortices, eds Peters A, Jones EG (Springer US, Boston, MA), pp 179–226.

16. Augustine JR (1996) Circuitry and functional aspects of the insular lobe in primates including humans. Brain Res Brain Res Rev 22(3):229–244.

17. Mesulam MM, Mufson EJ (1982) Insula of the old world monkey. III: Efferent cortical output and comments on function. J Comp Neurol 212(1):38–52.

18. Augustine JR (1985) The insular lobe in primates including humans. Neurol Res 7(1):2–10.

19. Kurth F, et al. (2010) Cytoarchitecture and probabilistic maps of the human posterior insular cortex. Cereb Cortex N Y N 1991 20(6):1448–1461.

20. Seeley WW, et al. (2012) Distinctive Neurons of the Anterior Cingulate and Frontoinsular Cortex: A Historical Perspective. Cereb Cortex 22(2):245–250.

21. von Economo C (1926) A new type of special cells of the cingulate and insular lobes. Z Ges Neurol Psychiatr 100:707–712.

22. Seeley WW, et al. (2007) Dissociable Intrinsic Connectivity Networks for Salience Processing and Executive Control. J Neurosci 27(9):2349–2356.

23. Critchley HD, Wiens S, Rotshtein P, Ohman A, Dolan RJ (2004) Neural systems supporting interoceptive awareness. Nat Neurosci 7(2):189–195.

24. Dosenbach NUF, et al. (2007) Distinct brain networks for adaptive and stable task control in humans. Proc Natl Acad Sci U S A 104(26):11073–11078.

25. Nimchinsky EA, et al. (1999) A neuronal morphologic type unique to humans and great apes. Proc Natl Acad Sci 96(9):5268–5273.

26. Deen B, Pitskel NB, Pelphrey KA (2011) Three systems of insular functional connectivity identified with cluster analysis. Cereb Cortex N Y N 1991 21(7):1498–1506.

27. Ryali S, Chen T, Padmanabhan A, Cai W, Menon V (2015) Development and validation of consensus clustering-based framework for brain segmentation using resting fMRI. J Neurosci Methods 240:128–140.

28. Chang LJ, Yarkoni T, Khaw MW, Sanfey AG (2013) Decoding the role of the insula in human cognition: functional parcellation and large-scale reverse inference. Cereb Cortex N Y N 1991 23(3):739–749.

29. Faillenot I, Heckemann RA, Frot M, Hammers A (2017) Macroanatomy and 3D probabilistic atlas of the human insula. NeuroImage 150:88–98.

30. Calamante F, Jeurissen B, Smith RE, Tournier J-D, Connelly A (2018) The role of wholebrain diffusion MRI as a tool for studying human in vivo cortical segregation based on a measure of neurite density: Diffusion MRI for Studying Cortical Segregation Based on Neurite Density. Magn Reson Med 79(5):2738–2744.

31. Latour LL, Svoboda K, Mitra PP, Sotak CH (1994) Time-dependent diffusion of water in a biological model system. Proc Natl Acad Sci 91(4):1229–1233.

32. Thornton JS, et al. (1997) Anisotropic water diffusion in white and gray matter of the neonatal piglet brain before and after transient hypoxia-ischaemia. Magn Reson Imaging 15(4):433–440.

33. Basser PJ, Pierpaoli C (1996) Microstructural and physiological features of tissues elucidated by quantitative-diffusion-tensor MRI. 111(3):209–219.

34. Mukherjee P, et al. (2002) Diffusion-Tensor MR Imaging of Gray and White Matter Development during Normal Human Brain Maturation. Am J Neuroradiol 23:1445–1456.

35. Ball G, et al. (2013) Development of cortical microstructure in the preterm human brain. Proc Natl Acad Sci U S A 110(23):9541–9546.

36. Fukutomi H, et al. (2018) Neurite imaging reveals microstructural variations in human cerebral cortical gray matter. NeuroImage. doi:10.1016/j.neuroimage.2018.02.017.

37. Aggarwal M, Nauen DW, Troncoso JC, Mori S (2015) Probing region-specific microstructure of human cortical areas using high angular and spatial resolution diffusion MRI. NeuroImage 105:198–207.

38. McNab JA, et al. (2013) Surface based analysis of diffusion orientation for identifying architectonic domains in the in vivo human cortex. NeuroImage 69:87–100.

39. Özarslan E, et al. (2013) Mean apparent propagator (MAP) MRI: a novel diffusion imaging method for mapping tissue microstructure. NeuroImage 78:16–32.

40. Fick RHJ, Wassermann D, Caruyer E, Deriche R (2016) MAPL: Tissue microstructure estimation using Laplacian-regularized MAP-MRI and its application to HCP data. NeuroImage 134:365–385.

41. Mitra PP, Latour LL, Kleinberg RL, Sotak CH (1995) Pulsed-field-gradient NMR measurements of restricted diffusion and the return-to-the-origin probability. J Magn Reson A 114(1):47–58.

42. Schwartz LM, Hürlimann MD, Dunn K-J, Mitra PP, Bergman DJ (1997) Restricted diffusion and the return to the origin probability at intermediate and long times. Phys Rev E 55(4):4225–4234.

43. Evrard HC, Logothetis NK, Bud Craig AD (2014) Modular architectonic organization of the insula in the macaque monkey: Architectonic organization of macaque insula. J Comp Neurol 522(1):64–97.

44. Evrard HC, Forro T, Logothetis NK (2012) Von Economo Neurons in the Anterior Insula of the Macaque Monkey. Neuron 74(3):482–489.

45. Kanti V Mardia PEJ (2000) Directional statistics (J. Wiley) Available at: http://gen.lib.rus.ec/book/index.php?md5=9CA4587ADE429BF78BD485D24346A7B1.

46. Taylor KS, Seminowicz DA, Davis KD (2009) Two systems of resting state connectivity between the insula and cingulate cortex. Hum Brain Mapp 30(9):2731–2745.

47. Cauda F, et al. (2012) Meta-analytic clustering of the insular cortex: characterizing the meta-analytic connectivity of the insula when involved in active tasks. NeuroImage 62(1):343–355.

48. Jelescu IO, Veraart J, Fieremans E, Novikov DS (2015) Degeneracy in model parameter estimation for multi-compartmental diffusion in neuronal tissue. 29(1):33–47.

49. Ascoli GA, Donohue DE, Halavi M (2007) NeuroMorpho.Org: A Central Resource for Neuronal Morphologies. J Neurosci 27(35):9247–9251.

50. Sotiropoulos SN, et al. (2013) Advances in diffusion MRI acquisition and processing in the Human Connectome Project. 80:125–143.

51. Nguyen DV, Li J-R, Grebenkov DS, Le Bihan D (2014) A finite elements method to solve the Bloch–Torrey equation applied to diffusion magnetic resonance imaging. J Comput Phys 263:283–302.

52. Ding SL, et al. (2016) Comprehensive cellular-resolution atlas of the adult human brain. J Comp Neurol 524(16):3127–3481.

53. Sridharan D, Levitin DJ, Menon V (2008) A critical role for the right fronto-insular cortex in switching between central-executive and default-mode networks. Proc Natl Acad Sci 105(34):12569–12574.

54. Roberts M, Hanaway J (1970) Atlas of the human brain in section (Lea & Febiger, Philadelphia).

55. Paxinos G, Huang XF, Toga AW (2000) The rhesus monkey brain in stereotaxic coordinates (Academic Press, San Diego, CA).

56. Bonthius DJ, Solodkin A, Van Hoesen GW (2005) Pathology of the Insular Cortex in Alzheimer Disease Depends on Cortical Architecture: J Neuropathol Exp Neurol 64(10):910–922.

57. Cai W, et al. (2016) Causal Interactions Within a Frontal-Cingulate-Parietal Network During Cognitive Control: Convergent Evidence from a Multisite-Multitask Investigation. Cereb Cortex N Y N 1991 26(5):2140–2153.

58. Ham T, Leff A, Boissezon X de, Joffe A, Sharp DJ (2013) Cognitive Control and the Salience Network: An Investigation of Error Processing and Effective Connectivity. J Neurosci 33(16):7091–7098.

59. Chen T, et al. (2015) Role of the anterior insular cortex in integrative causal signaling during multisensory auditory-visual attention. Eur J Neurosci 41(2):264–274.

60. Jilka SR, et al. (2014) Damage to the Salience Network and Interactions with the Default Mode Network. J Neurosci 34(33):10798–10807.

61. Craig AD (Bud) (2005) Forebrain emotional asymmetry: a neuroanatomical basis? TRENDS Cogn Sci 9(12):566–571.

62. Craig AD (2002) How do you feel? Interoception: the sense of the physiological condition of the body. Nat Rev Neurosci 3(8):655–666.

63. van den Heuvel MP, Mandl RCW, Kahn RS, Pol HEH (2009) Functionally Linked Resting-State Networks Reflect the Underlying Structural Connectivity Architecture of the Human Brain. 30(10):3127–3141.

64. Smith SM, et al. (2009) Correspondence of the brain’s functional architecture during activation and rest. Proc Natl Acad Sci 106(31):13040–13045.

65. Cauda F, et al. (2011) Functional connectivity of the insula in the resting brain. NeuroImage 55(1):8–23.

66. Vogt BA, Nimchinsky EA, Vogt LJ, Hof PR (1995) Human cingulate cortex: Surface features, flat maps, and cytoarchitecture. J Comp Neurol 359(3):490–506.

67. Chen T, Cai W, Ryali S, Supekar K, Menon V (2016) Distinct Global Brain Dynamics and Spatiotemporal Organization of the Salience Network. PLOS Biol 14(6):e1002469.

68. Clark L, et al. (2008) Differential effects of insular and ventromedial prefrontal cortex lesions on risky decision-making. Brain 131(5):1311–1322.

69. Gläscher J, et al. (2012) Lesion mapping of cognitive control and value-based decision making in the prefrontal cortex. Proc Natl Acad Sci:201206608.

70. Cai W, Chen T, Ide JS, Li C-SR, Menon V (2017) Dissociable Fronto-Operculum-Insula Control Signals for Anticipation and Detection of Inhibitory Sensory Cue. Cereb Cortex 27(8):4073–4082.

71. Taghia J, et al. (2018) Uncovering hidden brain state dynamics that regulate performance and decision-making during cognition. Nat Commun 9(1):2505.

72. Wen X, Liu Y, Yao L, Ding M (2013) Top-Down Regulation of Default Mode Activity in Spatial Visual Attention. J Neurosci 33(15):6444–6453.

73. Cai W, Chen T, Szegletes L, Supekar K, Menon V (2018) Aberrant Time-Varying Cross-Network Interactions in Children With Attention-Deficit/Hyperactivity Disorder and the Relation to Attention Deficits. Biol Psychiatry Cogn Neurosci Neuroimaging 3(3):263–273.

74. Uddin LQ, et al. (2013) Salience network-based classification and prediction of symptom severity in children with autism. JAMA Psychiatry 70(8):869–879.

75. Palaniyappan L, Simmonite M, White TP, Liddle EB, Liddle PF (2013) Neural primacy of the salience processing system in schizophrenia. Neuron 79(4):814–828.

76. Novikov DS, Jensen JH, Helpern JA, Fieremans E (2014) Revealing mesoscopic structural universality with diffusion. Proc Natl Acad Sci U S A 111(14):5088–5093.

77. Novikov DS, Kiselev VG, Jespersen SN (2018) On modeling. Magn Reson Med 9:413–422.

78. Allman JM, Watson KK, Tetreault NA, Hakeem AY (2005) Intuition and autism: a possible role for Von Economo neurons. Trends Cogn Sci 9(8):367–373.

79. Brüne M, et al. (2010) Von Economo neuron density in the anterior cingulate cortex is reduced in early onset schizophrenia. Acta Neuropathol (Berl) 119(6):771–778.

80. Seeley WW, et al. (2006) Early frontotemporal dementia targets neurons unique to apes and humans. Ann Neurol 60(6):660–667.

81. Santos M, et al. (2011) von Economo neurons in autism: A stereologic study of the frontoinsular cortex in children. Brain Res 1380:206–217.

82. Hotelling H (1992) Relations Between Two Sets of Variates. Breakthroughs in Statistics, Springer Series in Statistics. (Springer, New York, NY), pp 162–190.

83. Milham MP, et al. (2018) An Open Resource for Non-human Primate Imaging. Neuron 100(1):61–74.e2.

84. Seidlitz J, et al. (2018) A population MRI brain template and analysis tools for the macaque. NeuroImage 170:121–131.

85. Alnæs M, et al. (2015) The FEniCS Project Version 1.5. Arch Numer Softw 3(100). doi:10.11588/ans.2015.100.20553.

