## Supplementary Material for "Quantitative modeling links *in vivo* microstructural and macrofunctional organization of human and macaque insular cortex, and predicts cognitive control abilities"

**This PDF file includes:**

Supplementary Text

Supplementary Figures S1, S2

Supplementary Tables S1, S2, S3

Supplementary Text

Return to Origin Probability Density of Water Molecules as Measured by Diffusion MRI

The diffusion MRI signal characterizes the probability that water molecules within a voxel experience, on average, a net displacement after a given diffusion time. This enables the measurement of the probability that, after a diffusion time t, the water particles in a voxel are in the vicinity of their starting position, namely the Return-to-Origin probability density(1) (which in its normalized form we term RTOP). Even when the experimental conditions depart from the seminal hypotheses relating the dMRI signal attenuation with displacement probabilities(2), the RTOP has been shown, theoretically and experimentally to be an index closely related, in cellular tissue, to cellular shape and organization(1, 3). Furthermore, an advantage of the RTOP is that it is simple to estimate it by integrating the dMRI signal across a 3D volume. The unnormalized RTOP measurement is obtained from the dMRI signal $S$ as:

$P\left( t \right)=\frac{1}{S\left( 0,t \right)}\int_{\mathbb{R}^{3}} S\left( q,t \right)dq$.

In this expression, $P\left( t \right)$is the unnormalized RTOP at a given diffusion time $t$, and $q$ is the wavevector related to the usual diffusion b-value as $q=\sqrt{b}/t$, and the diffusion time is $t=\Delta-\delta/3$ where $\Delta$ and $\delta$ are the gradient pulse separation and length respectively.

Theoretical results enable us to link unnormalized RTOP measurements to characteristics of the environment where diffusion takes place(1, 3). Specifically, if water is diffusing freely, like in the ventricles^42,43^:

$$P\left( t \right)={(4\pi Dt)}^{-3}$$

With $D$the diffusion constant of cortical spinal fluid. For water diffusing within a microscopic compartment of volume V and surface S, at short diffusion times, unnormalized RTOP becomes

$P\left( t \right)=\left( 4\pi Dt \right)^{-3}\left( 1+\frac{\sqrt{\pi}}{2}\frac{S}{V}\left( Dt \right)^{\frac{1}{2}}-\left( \frac{1}{3}\left( \frac{1}{R_{1}}+\frac{1}{R_{2}} \right)+\frac{\rho}{D} \right)\frac{S}{V}Dt+O\left( {(Dt)}^{3/2} \right) \right)$.

$R_{1}$ and $R_{2}$ are the radii of curvature of the surface and $\rho$ is the surface relaxivity. The RTOP formulation can be simplified at long diffusion times: if the compartment is restricted along $d$ dimensions, unnormalized RTOP becomes an expression converging asymptotically to

$$P\left( t \right)\sim l^{d-3}\left( Dt \right)^{(d-3)/2}$$

with $l$ the average characteristic length scale of the compartment. For a sphere-type restriction, which can model a simplified cellular soma, there are no unrestricted dimensions, hence d=0 and $l^{3}\propto V$ is the soma’s internal volume, hence $P\left( t \right)\sim l^{-3}\propto V^{-1}$.

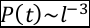

In practice, the ideal conditions for these unnormalized RTOP ansatzes are broken due to (i) the diffusion pulse length is finite; (ii) there is surface relaxivity; and, (iii) there is a finite maximum wavevector q that we are able to reach with the MRI equipment. Issue (i) has been shown to induce a distortion in the measurements making pores appear smaller(1, 3). Issue (ii) is generally mitigated by the normalization of $S(q;t)$ by $S\left( 0;t \right).$ Finally, issue (iii), restricts the size of the minimum observable compartment to $q^{-1}$, and to an underestimation of the real unnormalized RTOP value^42,43^.

Through this work, to render the unnormalized RTOP measurement, $P\left( t \right)$, comparable across different subjects we chose to use its normalized version(1), which we dub RTOP,

$R\left( t \right)=\left( 4\pi Dt \right)^{3}P(t)$,

obtaining the diffusion constant $D$ from the ventricles of each subject. RTOP, $R\left( t \right)$, which is dimensionless quantifies the relative enhancement of the unnormalized RTOP density, $P\left( t \right)$, with respect to free water diffusion due to restrictions(1, 3).

RTOP is sensitive to neuronal size

To assess that dMRI and RTOP, in our experimental conditions, is sensitive to the presence of Von Economo (VEN) and pyramidal neurons, we show analytically, through morphometric modelling simulations, and in an animal model, that the diffusion MRI signal at the acquisition parameters of clinical and pre-clinical MRI scanners, and specifically the HCP acquisition parameters, furthermore that its decay scales with the soma’s internal volume.

Our first simulation to assess the feasibility of obtaining a measurement related to neuronal size is purely theoretical. We analyzed the raw dMRI signal RTOP on VEN and pyramidal neurons. For this, we used Balinov et al’s(4) theoretical model to simulate dMRI acquisitions within a sphere to investigate whether the raw dMRI signal and RTOP are sensitive to the presence of large neurons. We assumed two samples of spherical volumes in agreement to Evrard et al’s(5) measurements of neuronal somas within the human fronto-insular cortical area: 15,347 ± 2,954 μm^3^ for VEN, and 5,226 ± 1,401 μm^3^ for pyramidal neurons. To simulate scenarios close to our human experimental data from HCP(6), we chose dMRI protocol using a gradient pulse length (δ) of 10.4 ms and separation (Δ) of 43.1 ms. For each soma we simulated two dMRI acquisitions, using 500 gradient strengths each, of G_max_ = 40 mT/m (b-­value = 509 s/mm^2^), corresponding to clinical scanners, and of G_max_ = 97 mT/m (b-value = 3000 s/mm^2^), corresponding to the pre-clinical scanner used in the HCP protocol.

Our results are shown in Figure S2. We found that at G_max_ = 40 mT/m the direction-averaged dMRI signal S_max_ at maximum gradient strength was -ln S_max_ = 0.46 ± 0.05, indicating a signal decay of 36 ± 3 % for large VEN-like neurons and -ln S_max_ = 0.20 ± 0.04, i.e. 20 ± 4 % decay for pyramidal-like neurons. This shows that at a clinically realistic(7) SNR of 20, or even at 10, both types of neurons will induce a detectable change in the dMRI signal. Furthermore, when looking the RTOP simulation results showing a RTOP of 1.51 ± 0.05 for large VENs and an RTOP of 1.74 ± 0.05 for pyramidals, the difference between these two measurements is an order of magnitude larger than the measurement variance, showing that the RTOP measurements are characteristic of each population.

These results are enhanced in the pre-clinical dMRI measurements available in the HCP dataset. When using a G_max_ = 97 mT/m, we obtained -ln S_max_ = 2.72 ± 0.31, characteristic of a decay of 93 ± 2 %, and for VEN and -ln S_max_ =1.29 ± 0.29, decay of 71 ± 9 %, for pyramidal. Thus, the dMRI signal, at G_max_ = 97 mT/m, is sensitive to neuronal size even at a signal-to-noise ratio of 1.

RTOP of 3.75 ± 0.50 for VENs and RTOP of 7.24 ± 1.13 for pyramidal neurons, showing a decrease in the differentiability but still a ratio of more than 3 standard deviations between them.

Furthermore, in the main text we have analyzed more realistic scenarios by simulating the HCP dMRI signal in realistic 3D digitally reconstructed models of fronto-insular human VEN and pyramidal neurons from NeuroMorpho.Org (8, 9), a Central Resource for Neuronal Morphologies. Furthermore, we used dMRI data from two macaque monkeys who were scanned using protocols similar to the HCP and shown that RTOP reflects known cytoarchitectural features of the primate insula using histological measurements of insular region cytoarchitectonics(5, 10, 11).

Taken together, these results show that, across multiple models and data acquisition protocols, including the one used in the HCP, the raw dMRI signal and RTOP are sensitive to the presence of VEN and pyramidal neurons.

Supplementary Figures

**Figure S1 Related to Figure 8. Analytical Simulation of Diffusion MRI Signal per neuron soma.** Neurons somas modelled as spheres with volumes reported by Evrard et al. On the left the dMRI signal separated by neuron type with the one standard deviation interval shaded, on the right histogram of normalized RTOP per neuron type. We show normalized RTOP for Gmax=97mT/m, as solid lines, corresponding to the maximum of our dataset and for Gmax=40mT/m, as dashed lines, corresponding to common clinical MRI scanners.

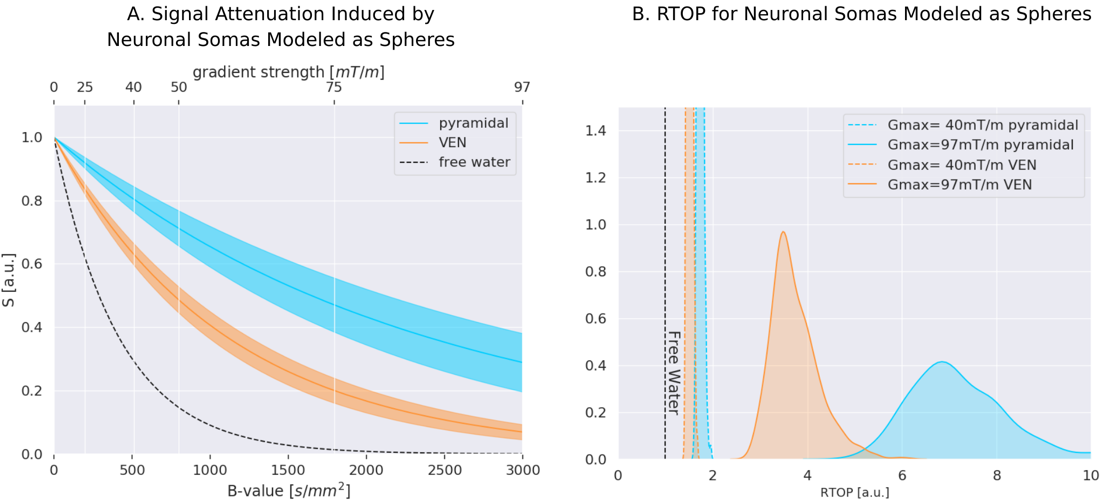

**Figure S2 Related to Figure 8.** **Simulated Diffusion MRI Signal per neuron type and section**. Simulations from three-dimensional neuron reconstructions issued from electronic microscopy. On the left the dMRI signal modulated by section partial volume and separated by neuron type and section, on the right the histogram of volume ratio between processes and soma for each neuron type.

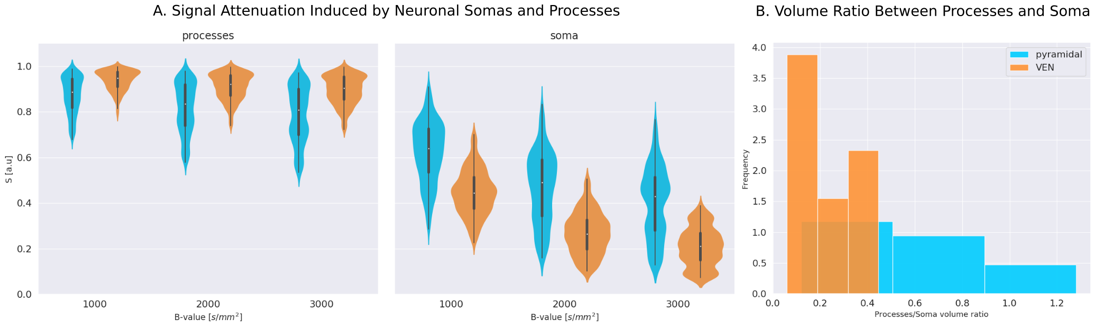

III. Supplementary Tables

**Table S1 Related to Figure 3.** **von Mises-Fisher statistics for the three insula subdivisions dAI, vAI, and PI.** For each subdivision, we computed the gradient’s main direction on each subject. The single subject 95% confidence interval (c.i.) summarizes the confidence intervals obtained across-subjects when computing their gradient’s mean direction. The population’s confidence interval shows the confidence interval for the mean direction computed from the subjects’ main directions. The Rayleigh statistic allows to reject the hypothesis of no preferential direction at the group level. The parcels with a p < 1e-10 where marked with an asterisk.

| Insula Subdivision | Single subject 95% c.i. (angle) | Group Level 95% c.i. (angle) | Rayleigh statistic |
| --- | --- | --- | --- |
| Left PI* | 0.24π +/- 0.10π | 0.02 π | 725 |
| Left dAI* | 0.24π +/- 0.11π | 0.02 π | 643 |
| Left vAI* | 0.28π +/- 0.10π | 0.04 π | 330 |
| Right PI* | 0.29π +/- 0.11π | 0.06 π | 136 |
| Right dAI* | 0.27π +/- 0.10π | 0.06 π | 485 |
| Right vAI | 0.30π +/- 0.10π | 0.08 π | 86 |

**Table S2 Related to Figure 6.** CCA Axis 1 weights.

| **RTOP measures** | **Weights** |
| --- | --- |
| rvAI | 0.17 |
| rPI | 0.09 |
| ldAI | 0.03 |
| lPI | 0.00 |
| lvAI | -0.13 |
| rdAI | -0.28 |
| **Behavior measures** | **Weights** |
| ListSort | 0.001 |
| Flanker | 0.001 |
| WM_Accuracy | 0.001 |
| CardSort | 0.001 |
| Gambling_Perc_Larger | 0.000 |
| WM_RT | 0.000 |
| Relational_RT | 0.000 |
| ProcSpeed | 0.000 |
| Gambling_RT_Larger | 0.000 |
| PicSeq | -0.002 |
| Relational_Accuracy | -0.003 |

Out-scanner behavioral measures:

“ListSort”, “Flanker”, “CardSort”, “ProcSpeed” and “Picseq” are standardized scores to evaluate individuals’ performance in the corresponding listing sorting, flanker, card sorting, processing speed and picture sequence tasks from the NIH Toolbox (<http://www.healthmeasures.net/explore-measurement-systems/nih-toolbox>).

In-scanner behavioral measures

“WM_Accuracy” stands for the accuracy in the n-back working memory task.

“WM_RT” stands for the reaction time in the n-back working memory task.

“Gambling_Perc_Larger” stands for the probability to select larger items in the gambling task.

“Gambling_RT_Larger” stands for the reaction time to select larger items in the gambling task.

“Relational_Accuracy” stands for the accuracy in the relational processing task.

“Relational_RT” stands for the reaction time in the relational processing task.

**Table S3 Related to Figure 7.** Comparison of microstructural properties of insular subdivisions in the macaque brain.

|  | **Ial versus the rest of the Insular Cortex Regions** | | | | | | | | | | | | |
| --- | --- | --- | --- | --- | --- | --- | --- | --- | --- | --- | --- | --- | --- |
|  | **X77** | | | | | | | **X181** | | | | | |
|  | **Left Hemisphere** | | | **Right Hemisphere** | | | | **Left Hemisphere** | | | **Right Hemisphere** | | |
|  | **t** | **p-value** | **df** | **t** | | **p-value** | **df** | **t** | **p-value** | **df** | **t** | **p-value** | **df** |
| Ig | -60.91 | < 0.001 | 4570 | -86.54 | | < 0.001 | 4872 | -76.73 | < 0.001 | 5040 | -76.19 | < 0.001 | 5650 |
| Id | -35.49 | < 0.001 | 4618 | -47.16 | | < 0.001 | 4948 | -52.25 | < 0.001 | 4584 | -51.91 | < 0.001 | 4729 |
| Iai | -55.04 | < 0.001 | 6573 | -51.60 | | < 0.001 | 8277 | -40.42 | < 0.001 | 6455 | -36.76 | < 0.001 | 6642 |
| Iapl | -25.47 | < 0.001 | 2402 | -31.76 | | < 0.001 | 2207 | -47.50 | < 0.001 | 3918 | -47.26 | < 0.001 | 4678 |
| Ia | 2.77 | 0.006 | 616 | -13.41 | | < 0.001 | 409 | 6.11 | < 0.001 | 733 | -2.89 | 0.004 | 825 |
| Iapm | 0.72 | 0.473 | 1121 | -17.35 | | < 0.001 | 1256 | 10.87 | < 0.001 | 1644 | -7.79 | < 0.001 | 1354 |
| Iam | -44.28 | < 0.001 | 3933 | -40.28 | | < 0.001 | 2446 | -49.24 | < 0.001 | 3770 | -59.67 | < 0.001 | 4266 |
|  | **Ig versus the rest of the Insular Cortex Regions** | | | | | | | | | | | | |
|  | **X77** | | | | | | | **X181** | | | | | |
|  | **Left Hemisphere** | | | **Right Hemisphere** | | | | **Left Hemisphere** | | | **Right Hemisphere** | | |
|  | **t** | **p-value** | **df** | **t** | **p-value** | | **df** | **t** | **p-value** | **df** | **t** | **p-value** | **df** |
| Ial | 60.91 | < 0.001 | 4570 | 86.53 | < 0.001 | | 4872 | 76.73 | < 0.001 | 5040 | 76.19 | < 0.001 | 5650 |
| Id | 49.61 | < 0.001 | 20062 | 75.61 | < 0.001 | | 20842 | 47.55 | < 0.001 | 18223 | 48.52 | < 0.001 | 17327 |
| Iai | -1.95 | 0.05 | 8209 | 18.68 | < 0.001 | | 6136 | 44.22 | < 0.001 | 9880 | 50.63 | < 0.001 | 10979 |
| Iapl | 12.02 | < 0.001 | 1413 | 16.88 | < 0.001 | | 1413 | 18.65 | < 0.001 | 1909 | 28.04 | < 0.001 | 2533 |
| Ia | 41.03 | < 0.001 | 376 | 20.31 | < 0.001 | | 328 | 63.15 | < 0.001 | 412 | 60.14 | < 0.001 | 424 |
| Iapm | 33.32 | < 0.001 | 773 | 26.28 | < 0.001 | | 881 | 72.68 | < 0.001 | 896 | 35.31 | < 0.001 | 959 |
| Iam | 4.75 | < 0.001 | 1703 | 14.20 | < 0.001 | | 1393 | 14.39 | < 0.001 | 1853 | 6.91 | < 0.001 | 2180 |

References

1.	Mitra PP, Latour LL, Kleinberg RL, Sotak CH (1995) Pulsed-field-gradient NMR measurements of restricted diffusion and the return-to-the-origin probability. *Journal of Magnetic Resonance, Series A* 114(1):47–58.

2.	Stejskal EO, Tanner JE (1965) Spin Diffusion Measurements: Spin Echoes in the Presence of a Time‐Dependent Field Gradient. *The Journal of Chemical Physics* 42(1):288–292.

3.	Schwartz LM, Hürlimann MD, Dunn K-J, Mitra PP, Bergman DJ (1997) Restricted diffusion and the return to the origin probability at intermediate and long times. *Phys Rev E* 55(4):4225–4234.

4.	Balinov B, Jonsson B, Linse P, Soderman O (1993) The NMR Self-Diffusion Method Applied to Restricted Diffusion. Simulation of Echo Attenuation from Molecules in Spheres and between Planes. 104(1):17–25.

5.	Evrard HC, Forro T, Logothetis NK (2012) Von Economo Neurons in the Anterior Insula of the Macaque Monkey. *Neuron* 74(3):482–489.

6.	Sotiropoulos SN, et al. (2013) Advances in diffusion MRI acquisition and processing in the Human Connectome Project. 80:125–143.

7.	Novikov DS, Kiselev VG, Jespersen SN (2018) On modeling. *Magnetic Resonance in Medicine* 9:413–422.

8.	Allman JM, et al. (2010) The von Economo neurons in frontoinsular and anterior cingulate cortex in great apes and humans. *Brain Struct Funct* 214:495–517.

9.	Ascoli GA, Donohue DE, Halavi M (2007) NeuroMorpho.Org: A Central Resource for Neuronal Morphologies. *J Neurosci* 27(35):9247–9251.

10.	Evrard HC, Logothetis NK, Bud Craig AD (2014) Modular architectonic organization of the insula in the macaque monkey: Architectonic organization of macaque insula. *Journal of Comparative Neurology* 522(1):64–97.

11.	Saleem KS, Logothetis N (2007) *A combined MRI and histology atlas of the rhesus monkey brain in stereotaxic coordinates* (Academic, London ; Burlington, MA).
